# Multimodal Imaging-Based Classification of PTSD Using Data-Driven Computational Approaches: A Multisite Big Data Study from the ENIGMA-PGC PTSD Consortium

**DOI:** 10.1101/2022.12.12.519838

**Authors:** Xi Zhu, Yoojean Kim, Orren Ravid, Xiaofu He, Benjamin Suarez-Jimenez, Sigal Zilcha-Mano, Amit Lazarov, Seonjoo Lee, Chadi G. Abdallah, Michael Angstadt, Christopher L. Averill, C. Lexi Baird, Lee A. Baugh, Jennifer U. Blackford, Jessica Bomyea, Steven E. Bruce, Richard A. Bryant, Zhihong Cao, Kyle Choi, Josh Cisler, Andrew S. Cotton, Judith K. Daniels, Nicholas D. Davenport, Richard J. Davidson, Michael D. DeBellis, Emily L. Dennis, Maria Densmore, Terri deRoon-Cassini, Seth G. Disner, Wissam El Hage, Amit Etkin, Negar Fani, Kelene A. Fercho, Jacklynn Fitzgerald, Gina L. Forster, Jessie L. Frijling, Elbert Geuze, Atilla Gonenc, Evan M. Gordon, Staci Gruber, Daniel W Grupe, Jeffrey P. Guenette, Courtney C. Haswell, Ryan J. Herringa, Julia Herzog, David Bernd Hofmann, Bobak Hosseini, Anna R. Hudson, Ashley A. Huggins, Jonathan C. Ipser, Neda Jahanshad, Meilin Jia-Richards, Tanja Jovanovic, Milissa L. Kaufman, Mitzy Kennis, Anthony King, Philipp Kinzel, Saskia B. J. Koch, Inga K. Koerte, Sheri M. Koopowitz, Mayuresh S. Korgaonkar, John H. Krystal, Ruth Lanius, Christine L. Larson, Lauren A. M. Lebois, Gen Li, Israel Liberzon, Guang Ming Lu, Yifeng Luo, Vincent A. Magnotta, Antje Manthey, Adi Maron-Katz, Geoffery May, Katie McLaughlin, Sven C. Mueller, Laura Nawijn, Steven M. Nelson, Richard W.J. Neufeld, Jack B Nitschke, Erin M. O’Leary, Bunmi O. Olatunji, Miranda Olff, Matthew Peverill, K. Luan Phan, Rongfeng Qi, Yann Quidé, Ivan Rektor, Kerry Ressler, Pavel Riha, Marisa Ross, Isabelle M. Rosso, Lauren E. Salminen, Kelly Sambrook, Christian Schmahl, Martha E. Shenton, Margaret Sheridan, Chiahao Shih, Maurizio Sicorello, Anika Sierk, Alan N. Simmons, Raluca M. Simons, Jeffrey S. Simons, Scott R. Sponheim, Murray B. Stein, Dan J. Stein, Jennifer S. Stevens, Thomas Straube, Delin Sun, Jean Théberge, Paul M. Thompson, Sophia I. Thomopoulos, Nic J.A. van der Wee, Steven J.A. van der Werff, Theo G. M. van Erp, Sanne J. H. van Rooij, Mirjam van Zuiden, Tim Varkevisser, Dick J. Veltman, Robert R.J.M. Vermeiren, Henrik Walter, Li Wang, Xin Wang, Carissa Weis, Sherry Winternitz, Hong Xie, Ye Zhu, Melanie Wall, Yuval Neria, Rajendra A. Morey

**Affiliations:** Department of Psychiatry, Columbia University Medical Center, New York, NY, USA; New York State Psychiatric Institute, New York, NY, USA; University of Rochester, Rochester, NY, USA; University of Haifa, Haifa, Israel; Tel-Aviv University, Tel Aviv, Israel; Baylor College of Medicine, Houston, TX, USA; Yale University School of Medicine, New Haven, CT, USA; University of Michigan, Ann Arbor, MI, USA; Duke University, Durham, NC, USA; Sanford School of Medicine, University of South Dakota, Vermillion, SD, USA; Munroe-Meyer Institute, University of Nebraska Medical Center, Omaha, NE, USA; University of California San Diego, La Jolla, CA, USA; Center for Trauma Recovery, Department of Psychological Sciences, University of Missouri-St. Louis, St. Louis, MO, USA; School of Psychology, University of New South Wales, Sydney, NSW, Australia; Department of Radiology, The Affiliated Yixing Hospital of Jiangsu University, Yixing, Jiangsu, China; Department of Psychiatry, University of Texas at Austin, Austin, TX, USA; University of Toledo, Toledo, OH, USA; University of Groningen, Groningen, The Netherlands; Minneapolis VA Health Care System, Minneapolis, MN, USA; University of Wisconsin-Madison, Madison, WI, USA; University of Utah School of Medicine, Salt Lake City, UT, USA; Departments of Psychology and Psychiatry, Neuroscience Program, Western University, London, ON, Canada; Department of Psychology, University of British Columbia, Okanagan, Kelowna, British Columbia, Canada; Medical College of Wisconsin, Milwaukee, WI, USA; UMR 1253, CIC 1415, University of Tours, CHRU de Tours, INSERM, France; Stanford University, Stanford, CA, USA; Emory University Department of Psychiatry and Behavioral Sciences, Atlanta, GA, USA; Civil Aerospace Medical Institute, US Federal Aviation Administration, Oklahoma City, OK, USA; Marquette University, Milwaukee, WI, USA; Brain Health Research Centre, Department of Anatomy, University of Otago, Dunedin, New Zealand; Department of Psychiatry, Amsterdam University Medical Centers, Academic Medical Center, University of Amsterdam, Amsterdam, The Netherlands; Brain Research and Innovation Centre, Ministry of Defence, Utrecht, The Netherlands; Cognitive and Clinical Neuroimaging Core, McLean Hospital, Belmont, MA, USA; Department of Radiology, Washington University School of Medicine, St. Louis, MO, USA; Division of Neuroradiology, Brigham and Women’s Hospital, Boston, MA, USA; School of Medicine and Public Health, University of Wisconsin-Madison, Madison, WI, USA; Heidelberg University, Heidelberg, Germany; University of Münster, Münster, Germany; University of Illinois at Chicago, Chicago, IL, USA; Ghent University, Ghent, Belgium; University of Cape Town, Cape Town, South Africa; Imaging Genetics Center, Mark and Mary Stevens Neuroimaging and Informatics Institute, Keck School of Medicine of the University of Southern California, Marina del Rey, CA, USA; Department of Psychology and Neuroscience, Baylor University, Waco, TX, USA; Wayne State University School of Medicine, Detroit, MI, USA; Division of Women’s Mental Health, McLean Hospital, Belmont, MA, USA; Department of Child and Adolescent Psychiatry, Psychosomatic and Psychotherapy, Ludwig Maximilian University of Munich, Munich, Germany; Psychiatry Neuroimaging Laboratory, Brigham and Women’s Hospital, Boston, MA, USA; Donders Institute for Brain, Cognition and Behavior, Centre for Cognitive Neuroimaging, Radboud University Nijmegen, Nijmegen, The Netherlands; Westmead Institute for Medical Research, Westmead, NSW, Australia; Department of Neuroscience, Western University, London, ON, Canada; University of Wisconsin-Milwaukee, Milwaukee, WI, USA; McLean Hospital, Belmont, MA, USA; Harvard Medical School, Boston, MA, USA; Institute of Psychology, Chinese Academy of Sciences, Beijing, China; Psychiatry and Behavioral Science, Texas A&M University Health Science Center, College Station, TX, USA; Department of Medical Imaging, Jinling Hospital, Medical School of Nanjing University, Nanjing, Jiangsu, China; University of Iowa, Iowa City, IA, USA; Charité Universitätsmedizin Berlin Campus Charite Mitte: Charite Universitatsmedizin Berlin, Berlin, Germany; VISN 17 Center of Excellence for Research on Returning War Veterans, Waco, TX, USA; Harvard University, Boston, MA, USA; Department of Psychiatry, Amsterdam University Medical Centers, VU University Medical Center, VU University, Amsterdam, The Netherlands; Department of Pediatrics, University of Minnesota, Minneapolis, MN, USA; Department of Psychology, Vanderbilt University, Nashville, TN, USA; University of Washington, Seattle, WA, USA; Department of Psychiatry and Behavioral Health, Ohio State University, Columbus, OH, USA; Neuroscience Research Australia, Randwick, NSW, Australia; Masaryk University, Brno, Czechia; Northwestern Neighborhood and Networks Initiative, Northwestern University Institute for Policy Research, Evanston, IL, USA; University of North Carolina at Chapel Hill, Chapel Hill, NC, USA; Center of Excellence for Stress and Mental Health, VA San Diego Healthcare System, San Diego, CA, USA; University of South Dakota, Vermillion, SD, USA; University of Minnesota, Minneapolis, MN, USA; Leiden University Medical Center, Leiden, The Netherlands; University of California, Irvine, Irvine, CA, USA; Department of Psychology, University of Chinese Academy of Sciences, Beijing, China

**Keywords:** posttraumatic stress disorder, multimodal MRI, machine learning, deep learning, classification

## Abstract

**Background:** Current clinical assessments of Posttraumatic stress disorder (PTSD) rely solely on subjective symptoms and experiences reported by the patient, rather than objective biomarkers of the illness. Recent advances in data-driven computational approaches have been helpful in devising tools to objectively diagnose psychiatric disorders. Here we aimed to classify individuals with PTSD versus controls using heterogeneous brain datasets from the ENIGMA-PGC PTSD Working group.

**Methods:** We analyzed brain MRI data from 3,527 structural-MRI; 2,502 resting state-fMRI; and 1,953 diffusion-MRI. First, we identified the brain features that best distinguish individuals with PTSD from controls (TEHC and HC) using traditional machine learning methods. Second, we assessed the utility of the denoising variational autoencoder (DVAE) and evaluated its classification performance. Third, we assessed the generalizability and reproducibility of both models using leave-one-site-out cross-validation procedure for each modality.

**Results:** We found lower performance in classifying PTSD vs. controls with data from over 20 sites (60% test AUC for s-MRI, 59% for rs-fMRI and 56% for d-MRI), as compared to other studies run on single-site data. The performance increased when classifying PTSD from HC without trauma history across all three modalities (75% AUC). The classification performance remained intact when applying the DVAE framework, which reduced the number of features. Finally, we found that the DVAE framework achieved better generalization to unseen datasets compared with the traditional machine learning frameworks, albeit performance was slightly above chance.

**Conclusion:** Our findings highlight the promise offered by machine learning methods for the diagnosis of patients with PTSD. The utility of brain biomarkers across three MRI modalities and the contribution of DVAE models for improving generalizability offers new insights into neural mechanisms involved in PTSD.

**Significance:**

⍰ Classifying PTSD from trauma-unexposed healthy controls (HC) using three imaging modalities performed well (∼75% AUC), but performance suffered markedly when classifying PTSD from trauma-exposed healthy controls (TEHC) using three imaging modalities (∼60% AUC).
⍰ Using deep learning for feature reduction (denoising variational auto-encoder; DVAE) dramatically reduced the number of features with no concomitant performance degradation.
⍰ Utilizing denoising variational autoencoder (DVAE) models improves generalizability across heterogeneous multi-site data compared with the traditional machine learning frameworks

## Introduction

Posttraumatic stress disorder (PTSD) is a prevalent and debilitating disorder, with a world-wide prevalence rate of 3.9% (1, 2). Current clinical assessments of PTSD rely solely on reported subjective experiences, overlooking objective biomarkers, which may lead to many cases of PTSD being undetected or misdiagnosed (3). Recent advances in computational power and data-driven computational approaches, especially supervised machine learning, have been helpful in devising tools to objectively diagnose psychiatric disorders (4-6). These approaches improve diagnosis by mining neuroimaging datasets, generating clinically relevant inferences at the individual level (7-9). In recent years, the number of supervised machine learning studies in translational neuroimaging has grown dramatically (10), but many challenges still remain. First, most extant studies are single-site studies of small homogeneous samples. Although efforts have been made to deal with overfitting (11, 12), single-site studies still tend to yield better performance than studies of larger samples, due to overfitting in the latter (13-15). Second, methodological differences across these studies (e.g., machine learning approaches, scanners, acquisition parameters, and data processing pipelines) limit the ability to directly compare their results. Third, most studies estimated classification performance via cross-validation (i.e., all samples are used in building the prediction model), without testing classification performance using independent yet-to-be-seen test data. For example, a recent review in depression has shown that only 4 of 66 studies evaluated classification performance using a holdout dataset, with all four containing less than 200 samples (9). However, for machine learning models to be useful in real-world clinical settings, predictive models need large samples that enable the evaluation of model performance on an unseen holdout dataset or independent cohorts. In PTSD, only a handful of studies exist, with none exploring the reproducibility of findings using multimodal brain imaging across multiple sites.

In addition to the above-described challenges, the selection of reliable and sensitive biomarkers to classify patients relative to controls is also crucial. In PTSD, most studies conduct group-level univariate analysis to identify PTSD-related biomarkers using one, and rarely two imaging modalities (16). No published studies thus far have explored three common imaging modalities of structural Magnetic Resonance Imaging (s-MRI), resting state functional MRI (rs-fMRI), and diffusion MRI (d-MRI), each tapping specific facets of structure or function to provide comprehensive information about the brain. S-MRI provides information on regional tissue volume of gray or white matter. In PTSD, structural abnormalities have been reported in the hippocampus, amygdala (17), prefrontal cortex, anterior cingulate cortex (18) and insula (19). Rs-fMRI measures the functional connectivity (FC) between brain regions. FC abnormalities in PTSD have been reported mainly in the default mode network (DMN), ventral attention network (VAN), executive control network (ECN) and salience network (SN) (20, 21). Finally, d-MRI provides information on white matter microstructure and the brain’s structural connectivity. White matter abnormalities in PTSD have been reported within the hippocampus, corpus callosum (22), cingulate gyrus (CG), uncinate fasciculus (23), and inferior fronto-occipital fasciculus (24, 25). However, as results from all three modalities are based on group-level analysis between PTSD and healthy controls (HC), or trauma exposed healthy controls (TEHC), it remains unclear whether PTSD can be discriminated at the single-subject level. Finally, most studies used only a single imaging modality among small samples (4, 26-28), limiting their broad-scale implications (4, 26, 29).

Recently, deep learning methods have received increasing attention in psychiatry because they are capable of learning subtle, latent patterns from high dimensional neuroimaging data. Deep learning methods have the potential to automatically diagnose different clinical disorders (30, 31), including PTSD (32), advancing the understanding of the neural basis of neuropsychiatric disorders (33). Of specific interest is autoencoder, which is a type of artificial neural network that seeks to learn the most efficient representations of the data at the individual level (34). Several neuroimaging studies show promising results for autoencoders in the classification of Alzheimer’s disease (35, 36), attention deficit hyperactivity disorder (ADHD) (37), autism spectrum disorder (ASD) (38), and schizophrenia (34, 39). Yet, the potential of autoencoders for multi-site and multimodal classification of PTSD remains unknown.

To address the gaps in knowledge, here we first used traditional machine learning approaches in large-scale multimodal heterogeneous datasets from the Enhancing Neuro-Imaging Genetics through Meta-Analysis (ENIGMA) and Psychiatric Genetics -(PGC) consortium PTSD working group. First, we assessed the utility of multimodal biomarkers including s-MRI, rs-fMRI, and d-MRI in classifying PTSD from healthy controls, both with and without trauma exposure, as previous research has suggested unique neural signatures associated with trauma-exposure that are not present in trauma-unexposed individuals (40, 41). To achieve this goal, we first identified the brain features that best distinguish PTSD from all non-PTSD controls. Next, we assessed the common and distinct neural features of PTSD versus controls with (TEHC) and without (HC) trauma exposure. Such information may provide valuable insight into underlying neural mechanisms in the pathophysiology of PTSD.

Second, we assessed the utility of deep learning models as a feature reduction method in improving classification performance. Neuroimaging studies usually make the predictive modeling task challenging because of the high dimensional feature set and relatively small sample size (42). Feature reduction methods can reduce feature dimensions to avoid overfitting, without losing important information needed for classification. Autoencoder approaches have an advantage over traditional feature reduction in suppressing noise from the input signal, leaving only a high-value representation of the input. Such an approach can automatically identify ways to transform raw imaging features into latent space variables, which are more suitable for machine learning algorithms, as well as capture the nonlinear representations of the input data. In this study, we built a denoising variational autoencoder (DVAE) for high dimensionality data reduction (43). The latent variables were used as new features and input into traditional machine learning approaches for classification. Instead of developing a system capable only of classifying individuals into patients and controls, we sought to capture the key feature information in the latent space using the DVAE model. We first trained the model using controls, and subsequently applied the model to data from PTSD patients. We aimed at retrieving the latent variables in PTSD patient data for capturing abnormal patterns of brain features (i.e., those that fall outside the normal range) (34).

Third, we assessed the generalizability and reproducibility of the classification model across heterogeneous datasets from multiple sites. The generalizability of machine learning to classify neuroimaging data is of great concern. Tremendous variability across studies inhibits the creation of a clear body of reliable knowledge from distinct studies (44). The ENIGMA-PGC consortium combines multimodal imaging and clinical data from multiple sites, enabling the development of models based on large samples. This offers an unprecedented opportunity for testing the generalizability and reproducibility of classification models to unseen datasets with vastly different characteristics compared to the sample used for model building. We evaluated generalizability across sites by assessing the classification performance for each site, and then by using Leave-One-Site-Out Cross-Validation (LOSOCV) to test how well the model generalized to independent cohorts.

## Methods

### 1. Dataset

Table 1 summarizes the descriptive information for each imaging modality. We analyzed brain MRI data from 6244 individuals (3,527 structural-MRI; 2,502 resting state-fMRI; and 1,953 diffusion-MRI). Of these 6244 individuals, 498 individuals had all 3 modalities, 736 had 2 and 5010 only had 1 modality. Inclusion and exclusion criteria for each cohort are summarized in *Supplemental Table 4*.

**Table 1:**
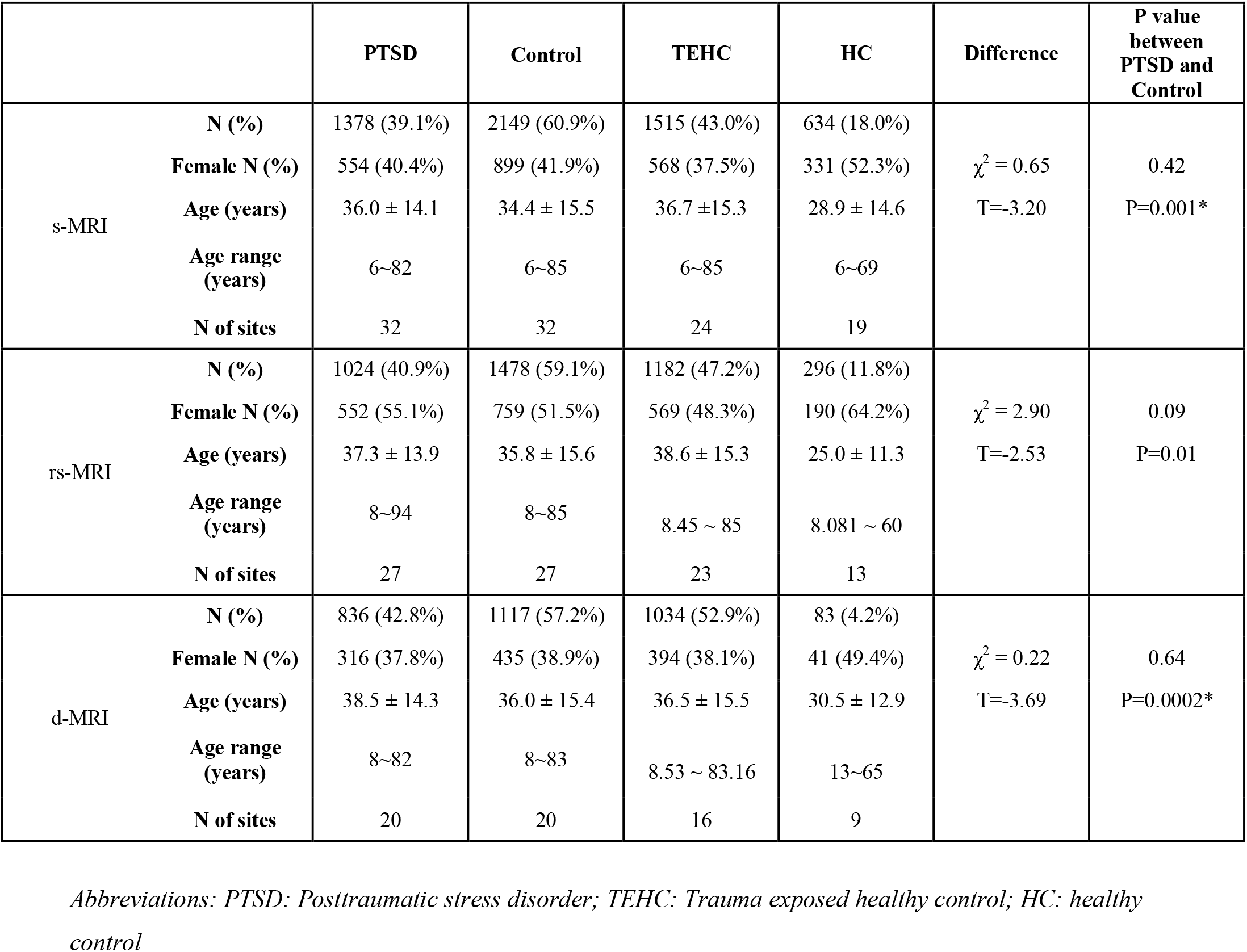
Demographics of PTSD and control groups across s-MRI (T1), rs-fMRI (RS), and d-MRI (DTI)

### 2. Image acquisition and processing

All imaging data were acquired at the contributing sites and processed with standardized protocols of the ENIGMA Consortium (45, 46). 96 cortical thickness and volume features were extracted from s-MRI, 10,878 pairwise connectivity measures were extracted from rs-fMRI based on the Power atlas (47), and 42 TBSS features were extracted from d-MRI. The overall set of imaging features used in this study are summarized in *Supplemental Figure 1 and supplemental Table 5∼7*.

### 3. Data Analysis

The overall analysis procedure is presented in **Figure 1**. For classification using traditional machine learning algorithms and DVAE, we used pooling methods for each modality, in which 70% of all sites’ data was used for cross-validation, and 30% of all sites’ data was used for holdout testing. For the generalization test, we first tested the classification performance for each site across all three modalities, and then used LOSOCV procedure for each modality. In addition, we tested the impact of site, age and sex on classification performance (*Supplemental Material*).

**Figure 1:**
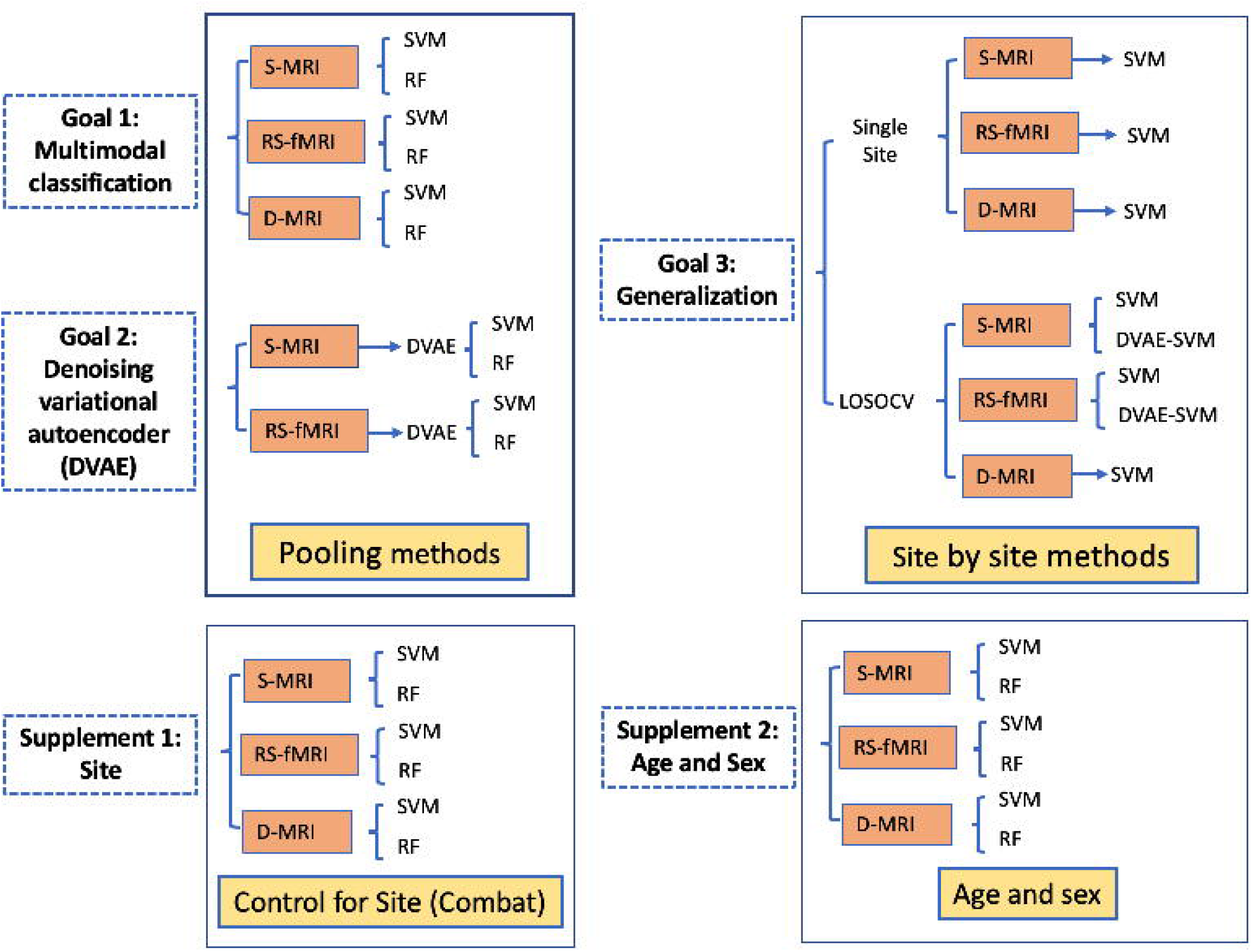
Overall analysis procedure. S-MRI: structural MRI; RS-fMRI: resting state fMRI; D-MRI: diffusion MRI; SVM: support vector machine; RF: random forest; DVAE: Denoising variational autoencoder.

#### Classification

We built three models for classifying PTSD relative to 1) all controls (HC and TEHC), 2) healthy controls with no trauma history (HC), and 3) those previously exposed to trauma who did not develop PTSD (TEHC) for each modality.

Support vector machine (SVM) and random forest (RF) algorithms were used for classification (*Supplemental Material*). Machine learning algorithms and Gridsearch were implemented in Python, and are available as part of the *scikit-learn* library. Our first task was to train classifiers that can differentiate patients with PTSD from control subjects in multi-site pooled data. In order to test the generalizability of the classifiers, we randomly split all imaging data into two subsets: 70% of the data was used for training and validation (cross validation), and the remaining 30% was used as a hold-out test data set. Brain features with 30% of missing data were dropped from further analysis (48). RobustScaler from the *scikit-learn* library was used to scale the data for each modality, and missing values were imputed with the mean of the training dataset. The same scaling method was applied to the test set (49). Gridsearch with stratified 10-fold cross-validation was used to select hyperparameters for both classifiers and to validate performance. Based on previous research (9), we used 10-fold cross validation, which generally provides better and more stable performance across different datasets, compared to Leave-One-Out Cross-Validation (LOOCV). To achieve an equal number of samples for each group, random under-sampling was applied to the imbalanced groups, with the under-sampling transform applied to the training dataset on each split of a repeated 10-fold cross validation. Classification performance was measured using standard metrics including accuracy, sensitivity, specificity, and area under the receiver operating characteristic curve (ROC-AUC), which summarizes sensitivity and specificity at different thresholds.

#### Denoising variational autoencoder (DVAE)

In our study, the feature dimension was very high for rs-fMRI data (148 ROIs, 10,878 ROI-to-ROI connectivity pathways), and relatively high for s-MRI data (96 regions). Researchers often use various feature reduction techniques for better performance and efficiency. Here, we implemented variational autoencoder models (VAE) using the PyTorch and TensorFlow libraries. The autoencoder consists of an encoder and a decoder. During the encoding phase, the VAE model learns a lower-dimensional representation (i.e. latent variables) for higher-dimensional data. The model was trained using rs-fMRI or s-MRI data from controls only. The resulting VAE model learned to encode healthy patterns from the input brain features into its latent representation. Later, the brain features from patients with PTSD were input into the same VAE model, and the latent variables were extracted as new features for machine learning analysis (**Figure 2**). The model architecture is described in S*upplemental Material: Methods*.

**Figure 2:**
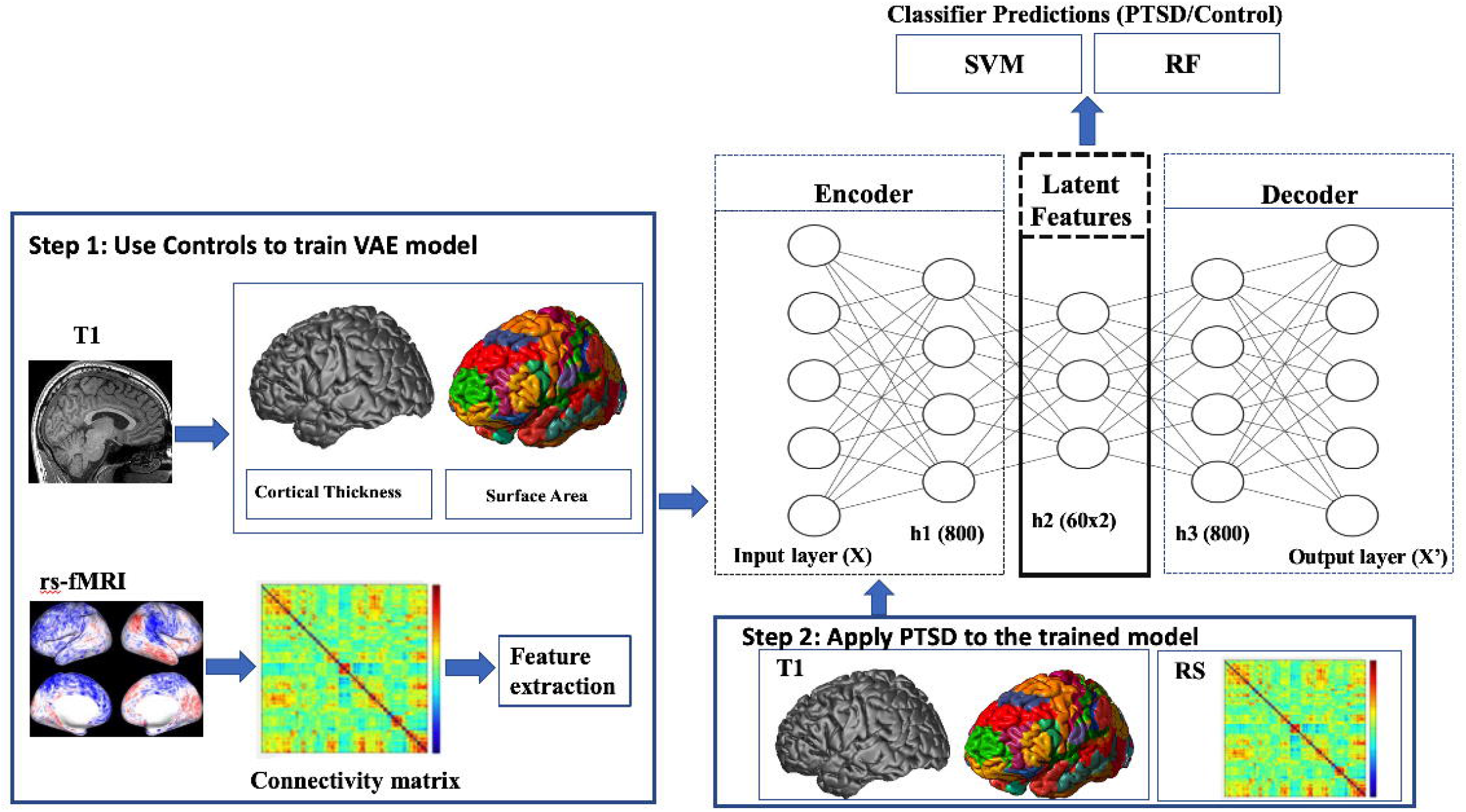
Denoising Variational Autoencoder analysis pipeline

## Results

### 1. Classification performance between PTSD and controls for each imaging modality using traditional SVM and RF

Figure 3 shows the CV AUC and test AUC for each modality using RF and SVM (s-MRI, rs-fMRI, d-MRI). The performance for RF was similar to SVM. Accuracy, Sensitivity and Specificity are reported in the *Supplemental Table 8 and Supplemental Figures 3-5*. First, our findings show that RF and SVM achieved similar performance when classifying PTSD from controls. Second, our models showed balanced CV AUC and test AUC, indicating that our models can generalize to an independent test set, which was not involved in model training, with no overfitting in these models. Third, all three modalities achieved comparable performance (using SVM: s-MRI: test AUC=60%, Cohen’s d=0.354; rs-fMRI: test AUC=59%, Cohen’s d=0.325; d-MRI: test AUC=56%, Cohen’s d=0.212). Among the three contrasts (PTSD vs. HC; PTSD vs. TEHC; PTSD vs Controls), the performance of classifying PTSD from HC was the best across all three modalities (SVM: s-MRI: test AUC=72%, Cohen’s d=0.82; rs-fMRI: test AUC=75%, Cohen’s d=0.948; d-MRI: test AUC=78%, Cohen’s d=1.09) (see *Supplemental Table 8*).

**Figure 3:**
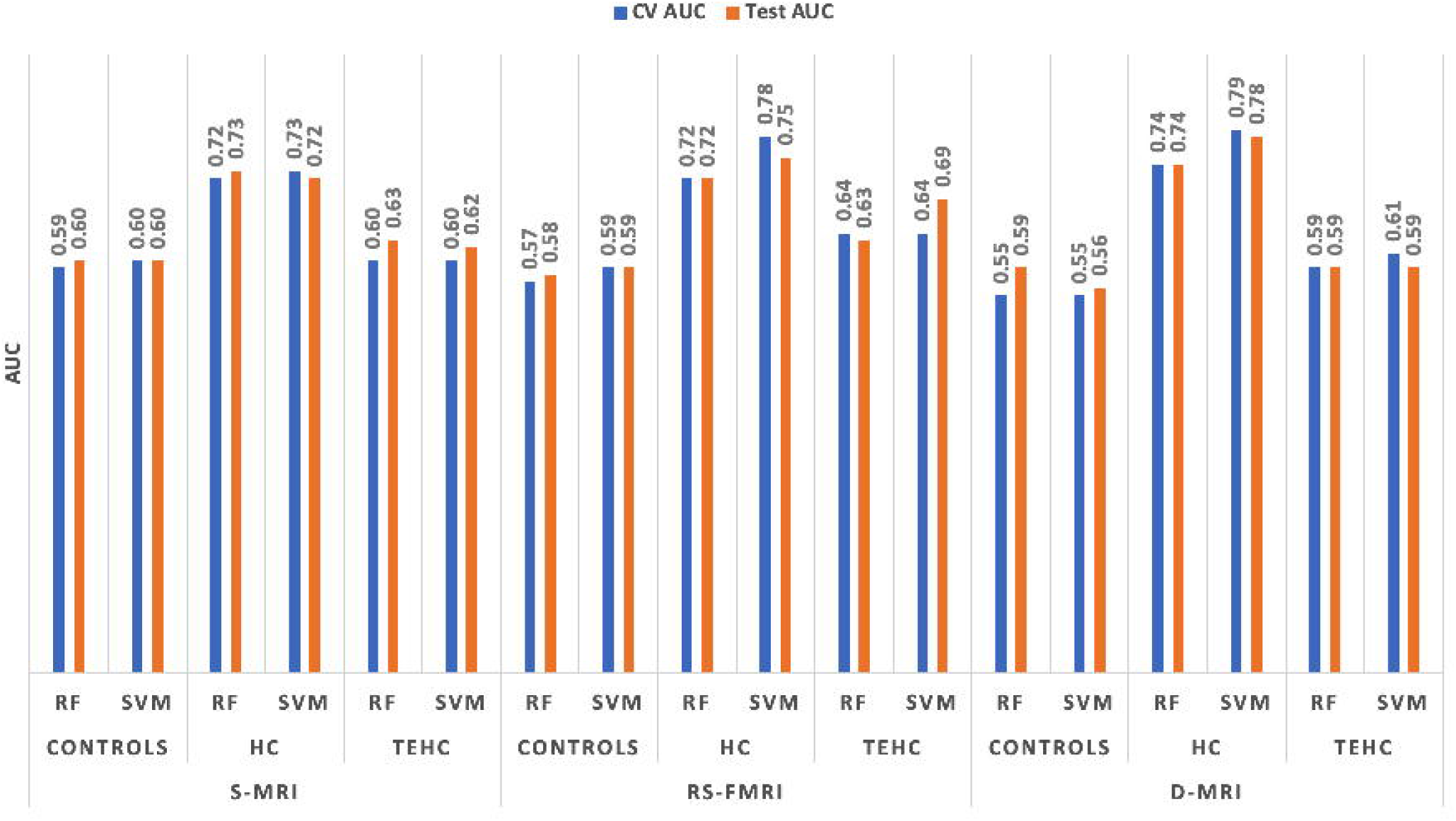
The overall classification performance (measured by cross validation AUC, and test AUC) between PTSD and all controls, between PTSD and HC, and between PTSD and TEHC, for S-MRI, rs-fMRI, and D-MRI

Some common and distinct features (*Supplemental Figure 6*) that differentiate PTSD from both HC and TEHC are presented in *Supplemental Results*.

### 2. Classification performance between PTSD and controls using deep learning framework

Applying DVAE to rs-fMRI data reduced the number of features (latent variables) to 10 based on a grid search, which were extracted from the original 10, 878 features. The performance of the model improved after feature reduction with DVAE (SVM: test AUC=62%, Cohen’s d=0.424). The performance of RF was the same as SVM (*Supplemental Figure 7*). We also applied DVAE to s-MRI data (feature size: 96; *Supplemental Table 5*), which reduced the features to 5 latent variables, and improved the performance after feature reduction with DVAE (SVM: test AUC=62%, Cohen’s d=0.424).

### 3. Generalization and reproducibility

#### 3.1 Assessing the classification performance for each site

S-MRI: The CV AUC in individual sites ranged from 0.42 to 0.82 using SVM. The average of individual site results yielded a CV AUC of 0.53 (std: 0.15), and CV accuracy of 0.64 (std: 0.11). Rs-fMRI: The CV AUC in individual sites ranged from 0.49 to 0.70 using SVM. The average of individual site results yielded a CV AUC of 0.56 (std: 0.07), and CV accuracy of 0.62 (std: 0.09). D-MRI: The CV AUC of individual sites ranged from 0.40 to 0.64 using SVM. The average of individual site results yielded a CV AUC of 0.52 (std: 0.076), and CV accuracy of 0.55 (std: 0.11) (**Figure 4**, and *Supplemental Table 9)*.

**Figure 4:**
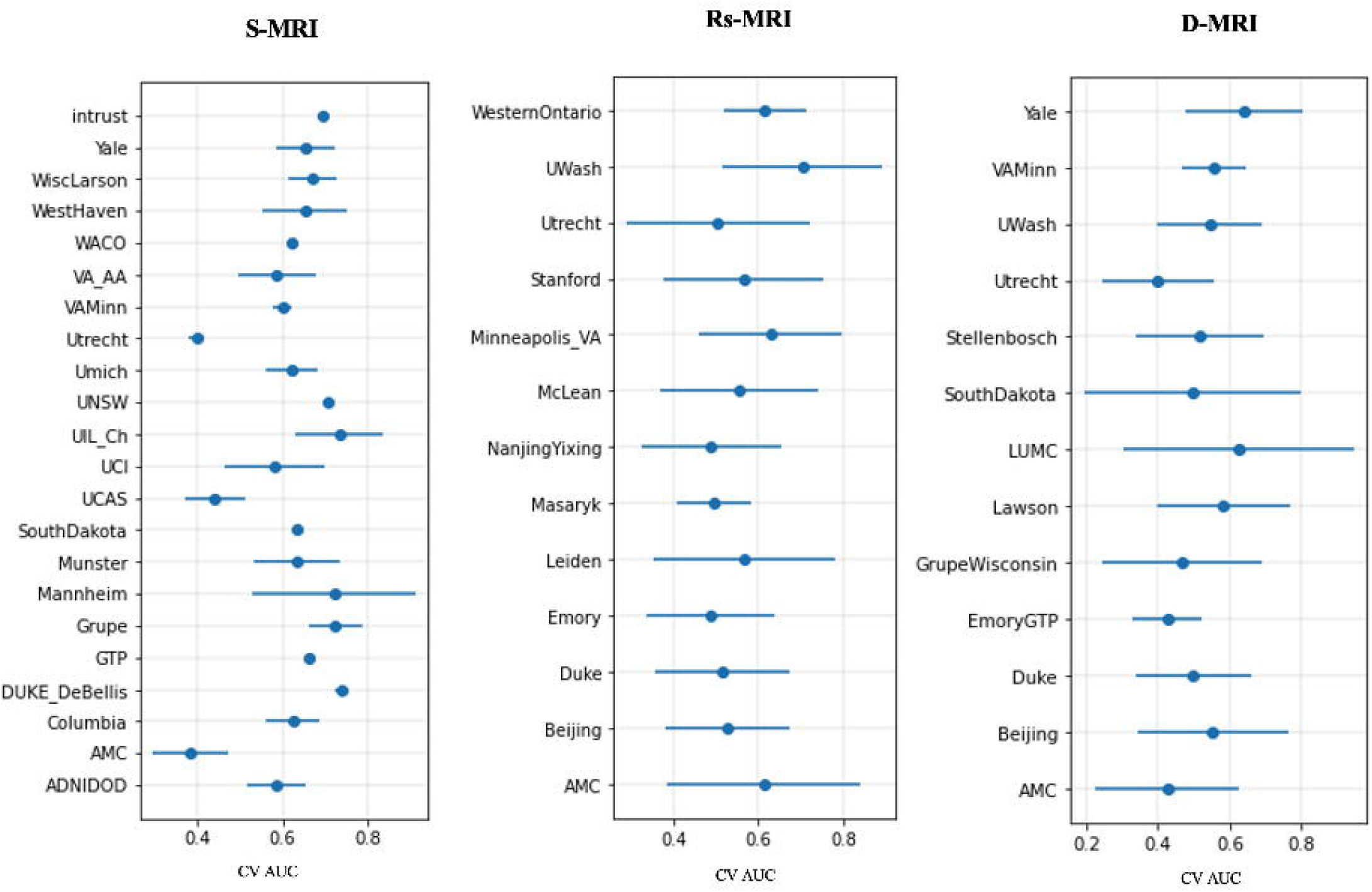
s-MRI, rs-fMRI, and d-MRI single site performance measured by CV AUC

We further assessed the correlation between the sample size at each individual site and the CV AUC. No significant correlation was found, for all three modalities.

#### 3.2 Leave one site out cross validation (LOSOCV)

The LOSOCV performance was compared with an aggregated pooling method (as in Results section 1). For all three MRI modalities (s-MRI, rs-fMRI, and d-MRI), LOSOCV provided chance level classification (s-MRI: Test AUC=56%, Cohen’s d=0.212; rs-fMRI: Test AUC=47%, Cohen’s d=0; d-MRI: test AUC=49%, Cohen’s d=0), and performed worse than the aggregate pooling method (*Supplemental Figure 9*). In s-MRI and rs-fMRI, we also compared LOSOCV and pooling methods using DVAE features, and assessed their generalizability. Only the DVAE achieved the same performance between LOSOCV and the pooling method, indicating a good generalization to unseen dataset using DVAE. Specifically, the LOSOCV method using SVM yielded an averaged CV AUC of 0.56 (std: 0.097) for s-MRI; an averaged CV AUC of 0.47 (std: 0.063) for rs-fMRI; and an averaged CV AUC of 0.49 (std: 0.080) for d-MRI (*Supplemental Figure 8*).

#### 3.3 Effects of site, age, and sex

We evaluated the impact of site by first harmonizing each site using ComBat (50), and then assessed the classification performance between PTSD and all controls using RF and SVM. The site harmonization did not impact the classification performance using RF, but the performance dropped using SVM (s-MRI: before: test AUC=60%, Cohen’s d=0.354; after: test AUC=52%, Cohen’s d=0.071; rs-fMRI: before: test AUC=59%, Cohen’s d=0.325; after: test AUC=46%, Cohen’s d=0; d-MRI: before: test AUC=56%, Cohen’s d=0.212; after: test AUC=52%, Cohen’s d=0.071) (*Supplemental Figure 10*).

We also evaluated the impact of age and sex on classification performance by including age and sex as features in the classification models. Age and sex did not impact the classification performance using either RF or SVM (*Supplemental Figure 11*).

## Discussion

The primary focus in the present study was to use machine learning techniques to create classifiers that leverage the complex multivariate patterns of structural and functional brain deficits. Specifically, we rigorously tested the classification performance on both cross-validation AUC and test AUC, in which a fully independent portion of the data was left out when selecting the model (both architectures and parameters). We found relatively poor classification performance in classifying PTSD vs. controls (60% test AUC for s-MRI, 59% for rs-fMRI and 56% for d-MRI using SVM), which is lower than top-performing studies conducted at a single site, ranging between 55.56% (13) and 97.1% (14) for rs-fMRI, and between 73% (26) and 80% (29) for studies focusing on multimodal biomarkers. Yet, single-site studies show poor generalization to independent datasets (51), suggesting that performance might be adversely affected by small sample sizes, high-dimensional features, and use of complex models with a large number of parameters. Good performance on training data, with poor performance on test data, suggests overfitting, as most machine-learning studies are evaluated only on the basis of cross validation. Therefore, while our accuracy is relatively low (13), the strength of our methods and sample size support the importance of our findings. Conversely, the present results are comparable with machine-learning studies using large scale imaging datasets in other psychiatric disorders based on s-MRI data – 65% accuracy when classifying MDD from HC (9); 65.2% accuracy in differentiating patients with bipolar disorder and from controls (45); and a CV AUC of 0.57-0.61 when classifying cases as OCD or controls (52). Exploring the utility of a DVAE, improved classification results emerged as compared to traditional ML approaches. The DVAE successfully reduced feature dimensions, e.g. reduced the rs-fMRI features from 10,878 features to 10 latent variables, without losing information important for classification (SVM test AUC=59%, Cohen’s d=0.325 using 10,878 features; test AUC=62%, Cohen’s d=0.424 using 10 latent variables). Thus, the present results have the potential to provide a baseline classification performance for PTSD when using large scale imaging datasets.

When considering HCs and TEHCs as separate control groups, our results yielded a markedly improved discrimination standard (test AUC in the range of 72% to 78%) across the three modalities, with the discrimination between PTSD and HC outperforming that of PTSD and TEHC. These findings are in line with previous studies showing greater similarity in underlying neural circuits between PTSD and TEHC participants (53, 54), than when comparing PTSD to HC with no trauma exposure.

Evaluating the generalizability by assessing the model performance for each site and each modality, showed that the classification AUC at the individual sites across all three imaging modalities ranged from 40% to 82% using SVM. However, such a wide range in the performance across individual sites is expected in large-scale multi-site studies, also shown in other disorders (45, 52), as samples are highly heterogeneous due to between-site differences (e.g., inclusion/exclusion criteria, demographic characteristics, clinical profiles, scanner used and scanning parameters, etc.). Furthermore, to avoid overfitting, we limited the scope of default parameters to SVM only, without hyperparameter tuning, which may have impacted the range of site performances compared to fine-tuning models using cross validation (45).

Our results also indicated that LOSOCV performed using traditional machine learning was worse than the performance using DVAE framework, as typically LOSOCV using traditional machine learning may result in large between-sample heterogeneity between training and test sets, resulting in roughly chance-level classification. Thus, imaging features do not provide strong biomarkers that enable generalization to new sites using traditional machine-learning methods. Previous studies have made an attempt at LOSOCV, yielding average accuracies of around 75.0% schizophrenia using s-MRI (55, 56), and an accuracy of 58.67% when assessed LOSOCV in bipolar disorder (45). These studies, however, used relatively small number of sites (3 to 5) for LOSOCV test, while we tested generalizability in 28 sites (s-MRI). Conversely, when extracting s-MRI and rs-fMRI features using DVAE models based on controls’ data only, the LOSOCV method achieved the same performance as the pooling method, demonstrating better generalizability using the DVAE framework. Importantly, the LOSOCV may be more significant in clinical practice because when multi-site data is used for model training, the final neuroimaging-based diagnostic classification models are much less site-specific, rendering them more generalizable.

Assessing the effect of *site* on classification performance showed that discrimination remained the same when using a random forest classifier, and dropped when using the SVM classifier, and after site harmonization with ComBat. Previous literature suggests that statistical harmonization methods developed to reduce data heterogeneity have the potential to improve accuracy, but at the cost of generalizability. Such approaches may compromise the train/test separation and introduce additional assumptions. Our findings demonstrate that DVAE may be able to capture differences across sites, and can be better generalized to new sites data. More importantly, the DVAE model does not require *a priori* knowledge of site information. Taken together, our findings support reproducibility of the DVAE across heterogeneous datasets from multiple sites. Testing generalizability, we also assessed the effect of *age* and *sex* on performance by adding them as features to the model (52), which did not affect classification performance. Neither did they emerge as informative features in classifying patients with PTSD from controls. Future studies should further assess the specific effects of *age* and *sex* on PTSD classification.

Several challenges still remain to be explored. First, combining biomarkers from different modalities with data fusion approaches is still in its infancy, and should be considered in future analyses to better detect potentially weak or latent effects hidden within high-dimensional datasets. Most deep-learning models are still being applied as black boxes, but serious efforts are underway to visualize latent variables and therefore improve the interpretability of results. Second, neuropsychiatric comorbidity was not consistently recorded across participating sites, so we could not evaluate it in the present study. Future studies should rectify this by also assessing comorbid conditions, exploring the underlying brain features that discriminate PTSD patients with and without comorbidity. Third, due to limited data of neurocognitive performance, we were not able to link emergent brain biomarkers to neurocognitive performance associated with the same brain circuits and/or regions. Fourth, while deep learning models typically give better performance than traditional machine learning, they are still perceived as black-box models, as they do not readily provide corresponding interpretations. However, deep learning need not be uninterpretable - as witnessed by the rapid expansion of methods for explainable deep learning (57, 58), which uses new forms of visualization and representations of model outcomes. Fifth, the deep learning models were trained using data from all controls, not HC; future studies could generate separate models using HC and TEHC, and further explore the difference in latent variables generated by different control groups (HC and TEHC). Lastly, although our study benefited from a large sample size and advanced analytics, its value in predicting disease progression and treatment response needs to be investigated by future studies.

Taken together, our findings highlight the promise offered by machine learning and deep learning methods in diagnosing patients with PTSD using multimodal brain imaging data. Our findings show that the control group used can heavily affect classification performance. We also demonstrate the possibility of improving generalizability using DVAE models, which may provide valuable insight into the neural mechanisms underlying the pathophysiology of PTSD.

## Acknowledgements

Dr. Zhu is supported by NIH K01MH122774 and by a NARSAD Young Investigator Grant from the Brain & Behavior Research Foundation 27040; Dr. Dennis is supported by NIH R61NS120249; Dr. Jahanshad is supported by NIH R01MH117601; Dr. Thompson is supported by NIH U54 EB020403; Dr. Fani is supported by NIH AT011267 and MH111671; Dr. Bomyea is suppprted by NIH R61MH127005 and CX001600; Dr. Lebois is supported by NIH K01MH118467; Dr. Daniels is supported by German Research Foundation DA 1222/4-1; Dr. Disner is supported by VA RR&D Award IK2RX002922; Dr. Bruce is supported by NIH K23 MH090366; Dr. Bryant is supported by National Health and Medical Research Council #1073041; Dr. Ross is supported by the NIH T32MH018931, F31MH122047 and T32GM007507; Dr. Cisler is supported by NIMH MH119132 and MH097784; Dr. Morey is supported by NIMH MH111671 and MH129832.

