## supplemental material for "Multimodal Imaging-Based Classification of PTSD Using Data-Driven Computational Approaches: A Multisite Big Data Study from the ENIGMA-PGC PTSD Consortium"

**Methods:**

**Clinical data:**

Depending on the cohort, current PTSD was diagnosed according to the Diagnostic and Statistical Manual of Mental Disorders (DSM) IV or V criteria, using the following standard instruments: Clinician-Administered PTSD Scale-IV (CAPS-IV), CAPS-5 (DSM-V), Structured Clinical Interview (SCID-IV) (DSM-IV), Mini International Neuropsychiatric Interview (MINI) 6.0.0 (3 cohorts, DSM-IV), PTSD Checklist (PCL)-4 (DSM-IV), PCL-5 (DSM-V), Davidson Trauma Scale (DTS) IV (1 cohort, DSM-IV), PTSD Symptom Scale (PSS) (DSM-IV), and Anxiety Disorders Interview Schedule (ADIS) (DSM-IV). All participating sites obtained approval from their local institutional review boards and ethics committees, and all study participants provided written informed consent.

**Image preprocessing:**

**S-MRI**: T1-weighted images were processed using the automated FreeSurfer processing stream to create individual subject thickness maps. The cortex of each hemisphere was parcellated into 34 cortical regions of interest (ROIs) using the Desikan–Killiany atlas (1). To match what we excluded for rs-fMRI data, 10 ROIs that are part of the motor or occipital lobes were removed from further analysis. The volume of an ROI was calculated by multiplying cortical thickness at each vertex in the ROI by the surface area across all vertices (2). ROI volumes and intracranial volume (ICV) were derived from subjects’ native spaces. Segmentations of gray and white matter and parcellations of ROIs were visually inspected using ENIGMA imaging quality control protocols (http://enigma.ini.usc.edu/protocols/). ROIs with segmentation or parcellation errors were excluded from the analysis. The final structural features included ROI cortical thicknesses (CT) and volumes for both left and right hemispheres, a total of 96 features (Supplemental Table 5).

R**s-fMRI:** Resting-state images were acquired at each site and preprocessed at a single location (Duke University). Preprocessing was implemented in ENIGMA HALFpipe workflow (https://github.com/HALFpipe/HALFpipe) based on fMRIPrep. Briefly, processing steps for T1w image include skull stripping, tissue segmentation, and spatial normalization to MNI space. Processing steps for functional images include motion correction using FSL MCFLIRT, slice time correction using AFNI 3dTshift for slice-timing correction, susceptibility distortion correction, and co-registration to the reference T1-weighted image using FSL FLIRT, and spatial normalization and warping to the template space using the MNI_2009 template. Each voxel was smoothed using signal from neighboring voxels with AFNI 3dBlurInMask followed by weighting by an isotropic Gaussian kernel. This method was repeated for each timepoint in the time series.

To ensure good quality of RS data, visual inspection was carried out on image registration, segmentation and brain extraction. To control confounding effects of motion artifact, several strategies were implemented: First, the top five aCompCor components were removed (3); second, frame-wise displacement (FD) was computed for each run, and subjects with more than 30% frames have high levels of gross motion were excluded (FD> 0.5 mm). Next, subjects with tSNR below 1.5 * IQR were excluded, and finally, subjects for whom more than 85% of independent component analysis (ICA) components classified as noise were further removed. The ROI-to-ROI functional connectivity was calculated by extracting the average time series extracted from each of the 264 ROIs regions defined by the Power atlas (4). A connectivity matrix between atlas regions was calculated using Pearson product moment correlation with PANDAS. We further reduced the number of features by only selecting 148 ROI regions that are part of known networks including default mode (DMN), ventral attention (VAN), frontal-parietal (FPN), salience (SN), subcortical (SCN), dorsal attention (DAN), cingulo-opercular networks (CON) (Supplemental Material Table 3) (5). The final functional connectivity feature set contained 10,878 measures.

**D-MRI:** DTI data were preprocessed following ENIGMA-DTI protocols and quality control procedures at (6). Processing steps include Eddy current correction, echo-planar imaging-induced distortion correction, motion correction, and tensor fitting. FA images generated from the estimated tensors were mapped to the ENIGMA DTI FA template and projected onto the skeleton FA template (FMRIB58_FA standard-space). FA values within ROIs were averaged within ROIs using JHU atlas for further analysis. Details and ROI abbreviations can be seen in *Supplemental Table 4*.

**Machine learning algorithms for classification:**

Support vector machine (SVM) is one of the most commonly used machine learning methods in all areas including psychiatric brain imaging studies. SVM algorithm aims to identify a hyperplane, or decision boundaries, that separates data points in a high dimensional feature space. Specifically, the hyperplane divides the space into two subspaces, each of which corresponds to one class, and it can be found by maximizing the margin between the two classes. After training, SVM classifiers can classify unlabeled test data based on which side of the hyperplane they lie (7, 8).

Random forest (RF) classifier is a popular ensemble learning algorithm that uses a combination of decision trees, where each tree casts a vote to classify the input vector. When building a RF classifier, the classification and regression trees learn from bootstrapped training data and are then tested on the out-of-bag samples, which allows for a wider diversity of trees. Random forest classifiers are known to be more robust, have higher accuracy compared to other classifiers, and run efficiently on large datasets (9).

**Calculating feature importance**: To find features that are most predictive of PTSD, we used a permutation-based feature-importance method on a RF classifier (10, 11). After choosing the best RF model, we permuted the values of each feature and recomputed the accuracy. Predictor importance was then described by the difference between the baseline accuracy of the classifier and the difference in accuracy after permuting the feature. This method, while slower to compute, is more robust than the *Gini importance* (GI) method, which is a more commonly used method to calculate feature importance.

**Denoising Variational autoencoder:**

*Model Architecture:* The autoencoder consists of an encoder and a decoder (S*upplemental Figure 2*). The encoder has one input layer, $x$, one hidden layer, $h1$, and an encoding layer, z. The decoder consists of one hidden layer, $h2$, and one output layer $\hat{x}$. The size of h1, h2, and z was varied depending on the modality used. For s-MRI, a size of h1 = h2 = 250 and size of z = 5 were chosen. For rs-fMRI, a size of h1 = h2 = 400 and a size of z = 10 was chosen. The sizes of the respective layers were chosen by performing a sparse grid search for each of the layers’ sizes independently and evaluating the performance of the model both with respect to the loss function and classification accuracy (12).

The encoding layer *z* is referred to as the *latent space* of the model. This layer stores the model’s reduced feature representation of the input data. In a general VAE framework, the features of the latent space *z*, referred to as *latent variables*, are drawn from Gaussian distributions determined by learned parameters ($\mu, log(\sigma^{2})$). These Gaussian distributions comprise an estimated distribution $q(z|x)$ to approximate the true underlying prior distribution $p(z)$. Once the encoded representation $z$ is sampled, the values are reparameterized and fed into the decoder network. The decoder network then tries to reproduce the input using the reparameterized encoded data. The activation function for the layers was chosen as scaled exponential linear units (SELU) (13).

*Loss Function of Model:* For an autoencoder, the loss is usually determined solely by $L=MSE(x,\hat{x})$ where MSE is the mean squared error loss. This makes the autoencoder’s sole objective to maximize its reconstruction accuracy. For a VAE the Kullback-Leibler Divergence (D_KL_) is added to the loss function. The D_KL_ term is used to determine how much $q(z|x)$ and $p(z)$ differ. This constrains the way in which the parameters for the Gaussian distributions are updated and regularizes the latent space. Thus for a VAE, the loss function is generally

$L=MSE\left( x,\hat{x} \right)+D_{KL}(q\left( x \right)||p(z))$.

For $p(z)$, usually the unit Gaussian or $N(0,1)$ is chosen. This was the choice for our model as well.

*Training of the DVAE model:* During training, the model was only fed the data of combined control participants (HC and TEHC). Our intent was that the model would first learn the features representing salient aspects of healthy brain function and use the same features to represent PTSD. Prior to feeding the data to the model, the data was standardized by median and interquartile range. Additionally, a Gaussian noise was applied with a mean of 0 and a standard deviation of 0.1 to the input data. Our goal was to induce the model to learn to find more robust features of the data that are tolerant to noise and thus be able to reconstruct the noiseless data while being fed noisy data. This was the *denoising* aspect of the DVAE. The samples were then split into a training (70%) and test (30%) set. Then 20% of the training set was set aside for validation. Each epoch, the samples were fed as mini batches of size 128 to the network. L2 regularization (regularization parameter = 0.1) was applied to penalize high values of the network’s weights and to avoid overfitting. The model was then trained for 500 epochs.

Convergence was measured by evaluating the *per-feature loss*, which we defined as $L/n$ where n is the number of features of the input. This was done so as to be able to roughly compare the loss for models from different modalities, as each had a different number of features.

*Encoding and Classification:* After training the model, we used it to encode and reconstruct the data from both control subjects and the individuals with PTSD. For each subject, the values for their latent distributions, $\mu, log(\sigma^{2})$, were computed and extracted. Second, we compared the performance of the encoded features to the original features using the SVM and RF classifiers.

*Measuring the Contribution of Latent Variables:* The contribution of each latent variable was evaluated by varying each latent variable independently. As each latent variable was changed, a new latent representation z was generated. The previous value and the current value of z were both fed into the decoder to produce a reconstruction and the absolute value of their difference was computed. This enabled the visualization of the contribution of each latent variable at its current value. Since the latent distributions for each subject looked different, this method allowed us to determine which features of the input were different for each subject. A similar analysis was performed on the averaged distributions for all controls and PTSD individuals, respectively, to examine how the model differentiates between them.

**Generalization:**

Single Site: We evaluated the same SVM parameterization used in previous analyses on each site’s data, to shed light on its replicability. However, this method requires each site's sample size to be large enough to appropriately fit a machine-learning model. Thus, we only included sites that had more than 20 subjects in each group (PTSD and all control). For sites that have imbalanced samples, a down sampling approach was used to have a distributed sample across the two groups. To maximize generalizability and avoid overfitting, we applied the SVM for each site using the default parameters (C = 1), without grid search for optimal parameters, or feature reduction and selection. This method is stratified insofar as the proportion of cases and controls (in respective folds) is similar in both training and validation sets. The SVM model was trained and evaluated using a 10-fold cross validation, and predictive performance was evaluated on the data from the held-out site.

Leave-one-site-out cross-validation (LOSOCV): We evaluated the same SVM parameterization used in all other analyses on a LOSOCV procedure. Sites with sample size greater than 20 in each group were included in this analysis. In each fold of cross-validation, one hold-out site was completely left out from the training partition, which was further randomly partitioned into 10 folds for cross validation. Predictive performance was evaluated on the data from the hold-out site. The goal of this procedure was to assess the generalizability of the classifier to a totally independent data set that was sampled from a different sample and scanner.

ComBat: For a large multi-site study, it is important to consider whether a classifier can well generalize to new data coming from a different scanner or country. We used the ComBat method (14) to remove the site-specific information from the data and to test the generalizability of our classifier. The ComBat method models each imaging measurements as a combination of three parts: variation of Y captured by the covariates such as age and sex, mean differences across sites, and the error term that follow varying normal distributions at different sites. Then the ComBat-transformed data can be achieved by removing these additive and multiplicative effects due to site differences.

Biologically relevant covariates: To evaluate the contribution of confounding factors such as age and sex on the classification performance, we included age and sex as features and tested the impact on the overall performance.

Code is available on GitHub:

<https://github.com/ColumbiaNeriaLab/MultimodalPTSDClassification>

**Results:**

**Top features identified that distinguish PTSD and controls for each modality using traditional SVM and RF**

Some common features (*Supplemental Figure 6*) that differentiate PTSD from both HC and TEHC were detected in 1) s-MRI features include brain regions within the default mode network (DMN [parahippocampal, supramarginal, precuneus, inferior parietal, lateral temporal cortex]), and within the prefrontal cortex (inferior frontal gyrus, rostral middle frontal); 2) rs-MRI features included DMN-SN, FPN-SN, DMN-FPN, DMN-VAN, FPN-DAN, DMN-DAN, and within DMN connectivity; and 3) d-MRI features included the cingulum (hippocampal portion; CGH), superior *corona radiata* (SCR), fornix (crus)/stria terminalis (FX_ST), anterior limb of internal capsule (ALIC), posterior limb of internal capsule (PLIC), uncinate fasciculus (UNC), CST, SFO, SLF, EC and SS. Other informative features that only differentiate PTSD from HC were detected in 1) s-MRI features included within the salience network (SN) (ACC and insula); 2) rs-fMRI features included CO-SC, CO-SN, and within SC; and 3) d-MRI features included ACR, TAP, BCC, RLIC. However, these features were not informative in distinguishing PTSD from TEHC. The most informative features from classifying PTSD from TEHC include 1) s-MRI of prefrontal cortex (orbitofrontal cortex, superior frontal cortex); 2) rs-fMRI connectivity between FPN-SC, and CO-DMN; and 3) d-MRI of internal capsule (IC), external capsule (EC), GCC, PTR, CGC, and SCC.

**Supplemental Figures:**

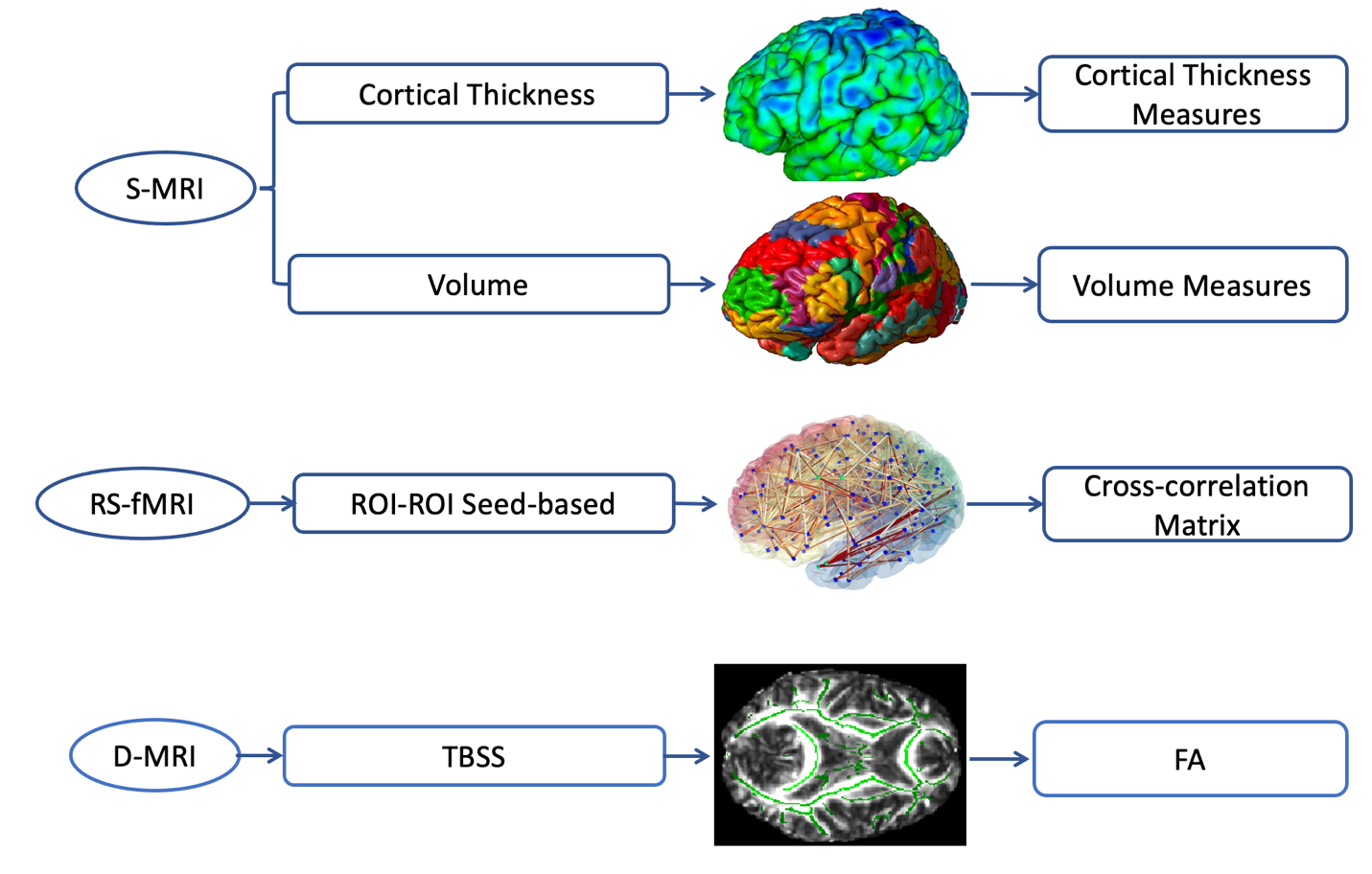

*Supplemental Figure 1: Brain features from structural MRI (s-MRI), resting state fMRI (rs-fMRI), and DTI (d-MRI) used in this study*

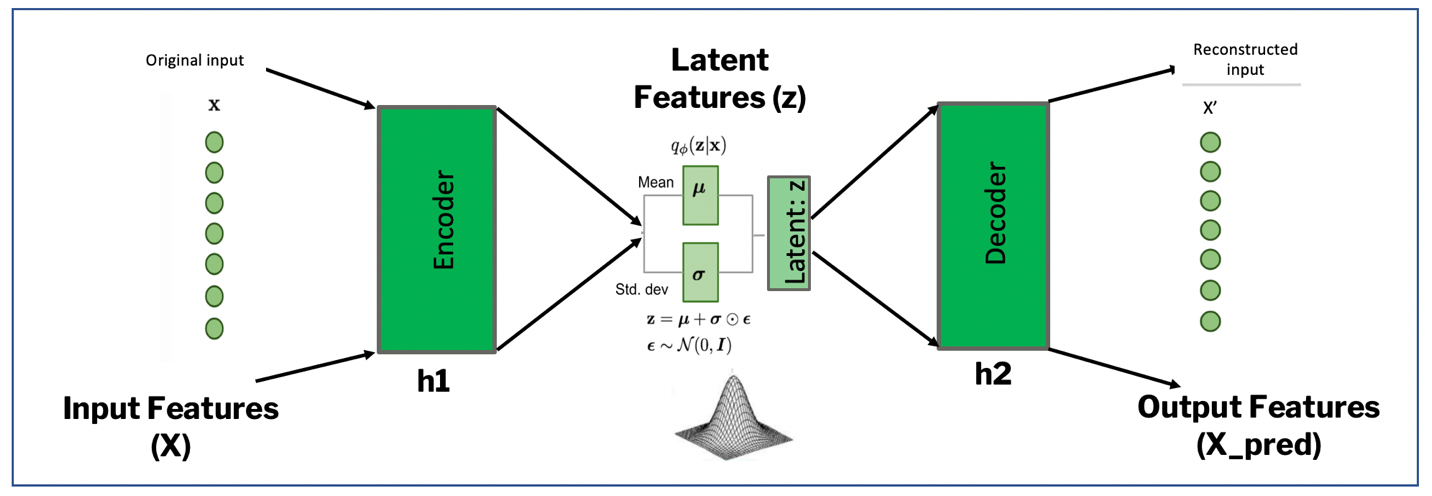

*Supplemental Figure 2: Denoising Variational Autoencoder model*

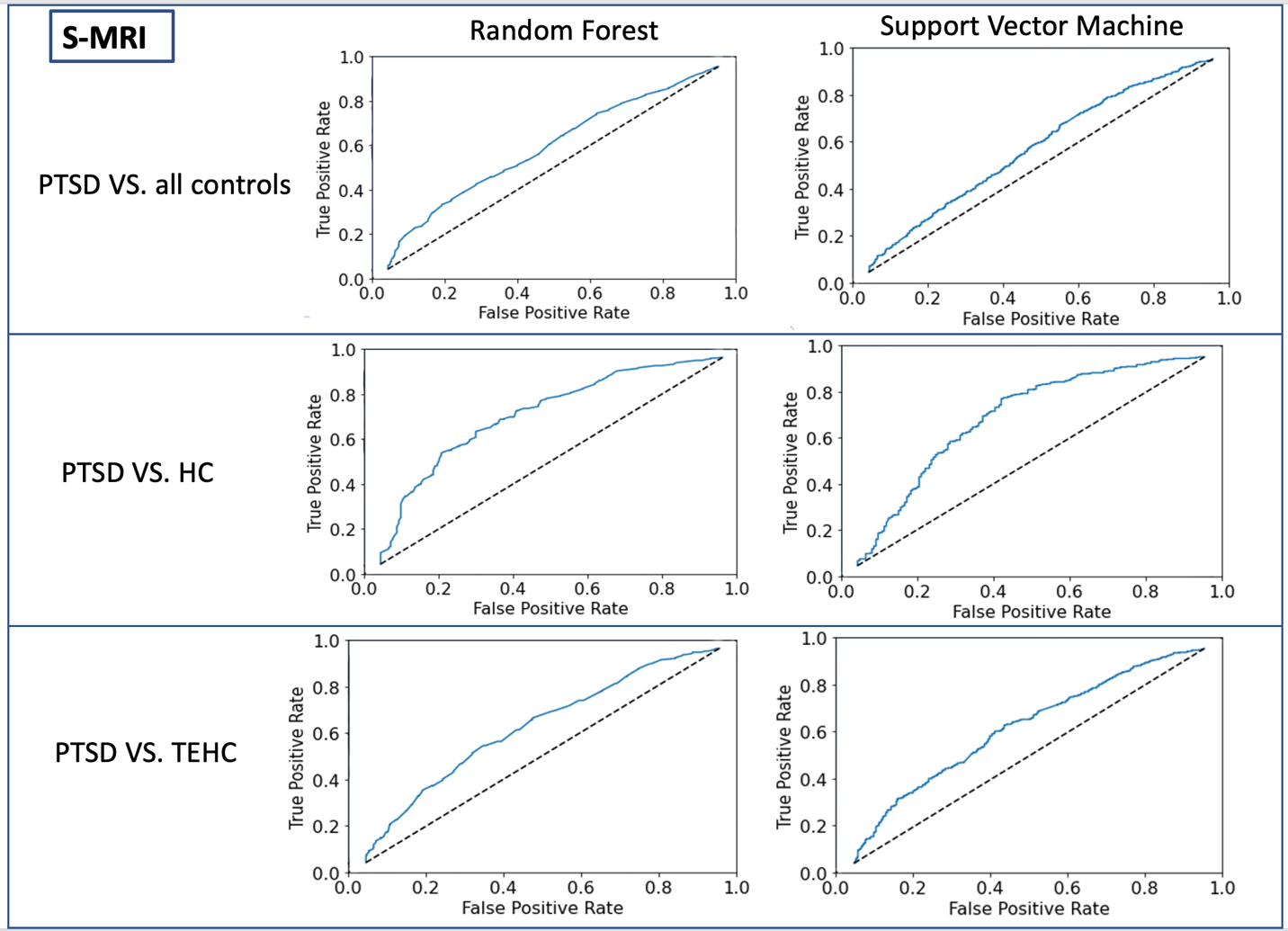

*Supplemental Figure 3: The classification performance using s-MRI measured by test ROC between PTSD and all controls (HC+TEHC), (top two figures, left: RF model, right: SVM model), between PTSD and HC (middle two figures, left: RF model, right: SVM model), and between PTSD and TEHC (bottom two figures, left: RF model, right: SVM model)*

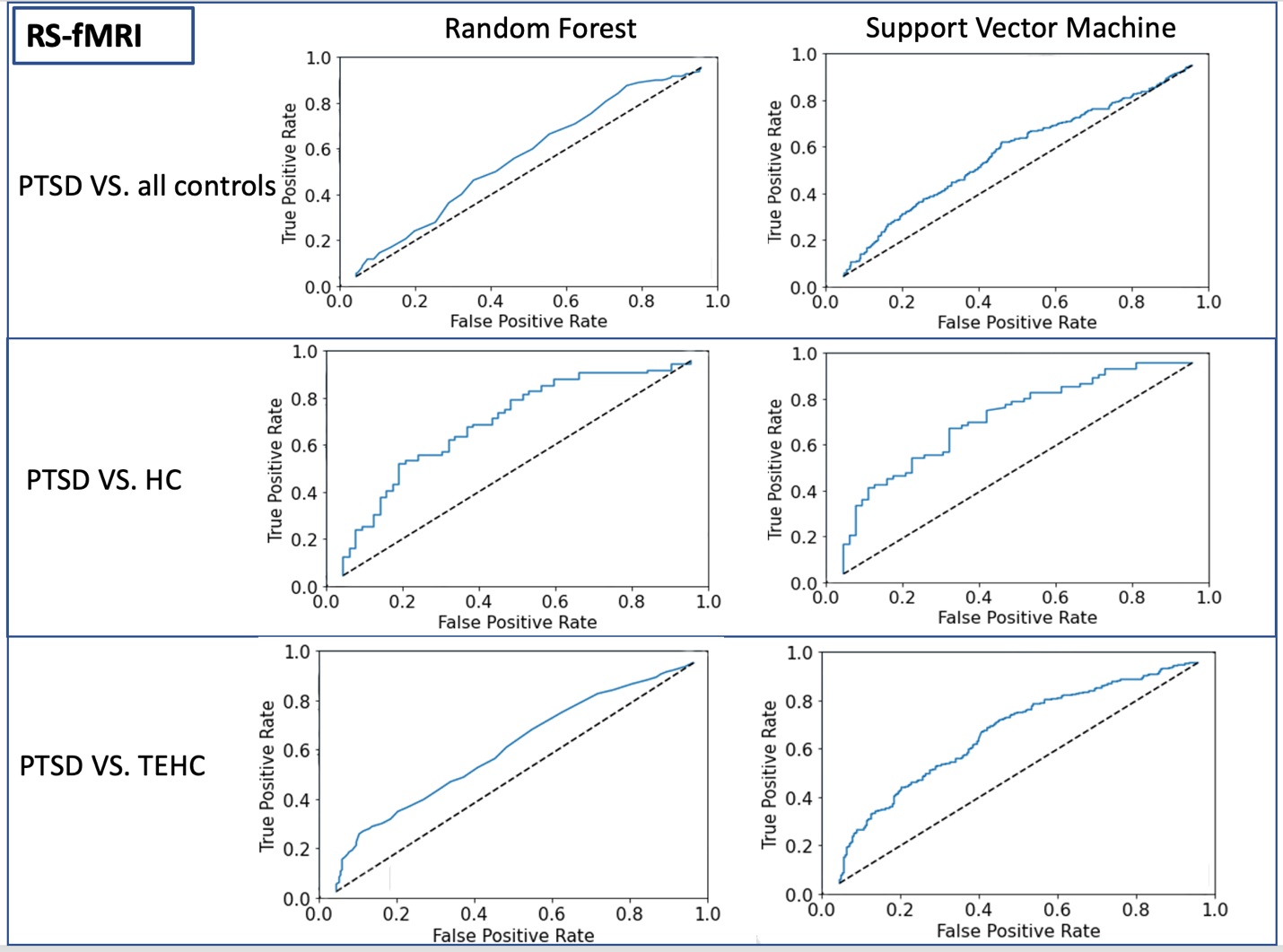

*Supplemental Figure 4: The classification performance using rs-MRI measured by test ROC between PTSD and all controls (HC+TEHC), (top two figures, left: RF model, right: SVM model), between PTSD and HC (middle two figures, left: RF model, right: SVM model), and between PTSD and TEHC (bottom two figures, left: RF model, right: SVM model)*

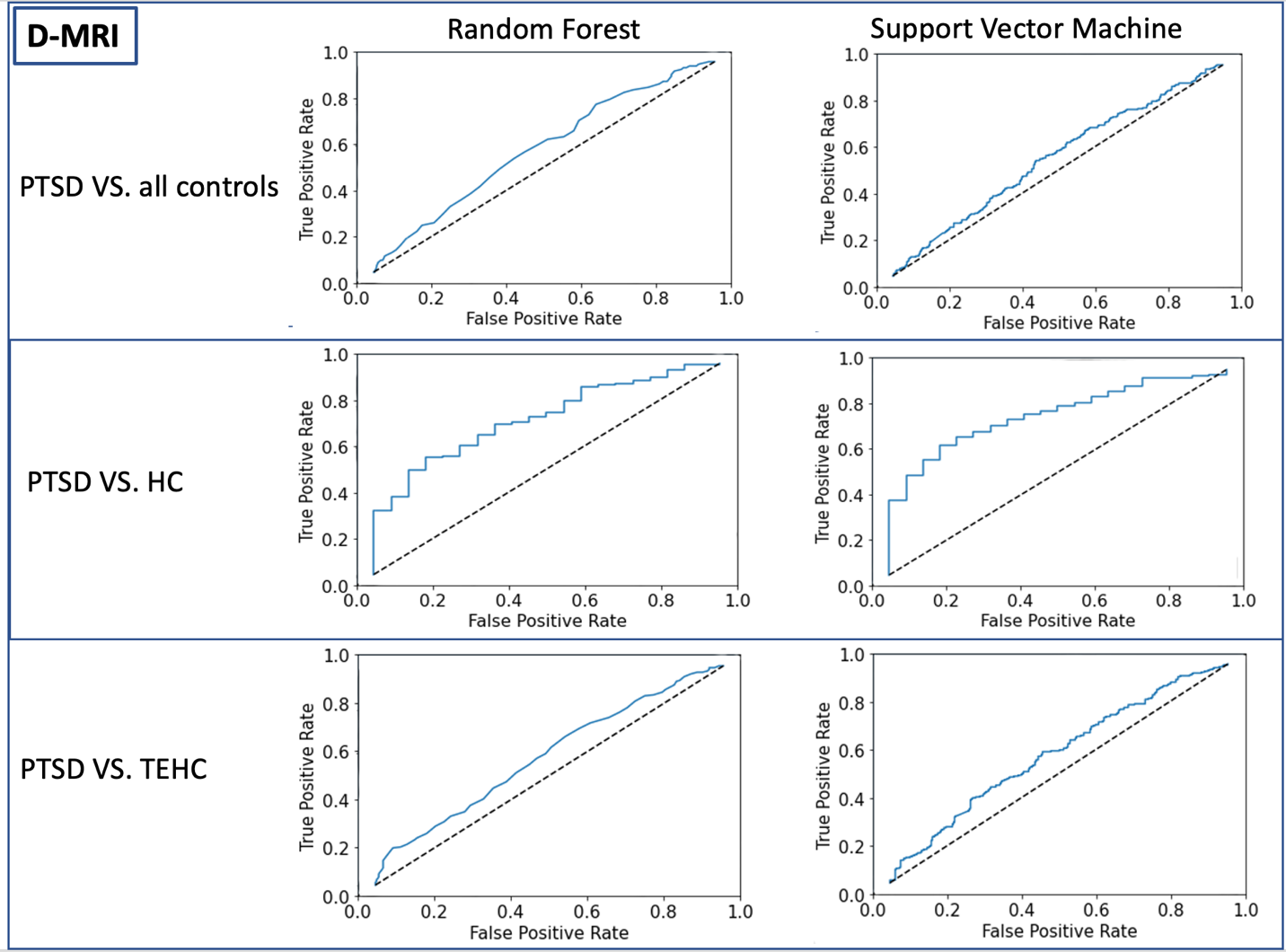

*Supplemental Figure 5: The classification performance using d-MRI measured by test ROC between PTSD and all controls (HC+TEHC), (top two figures, left: RF model, right: SVM model), between PTSD and HC (middle two figures, left: RF model, right: SVM model), and between PTSD and TEHC (bottom two figures, left: RF model, right: SVM model)*

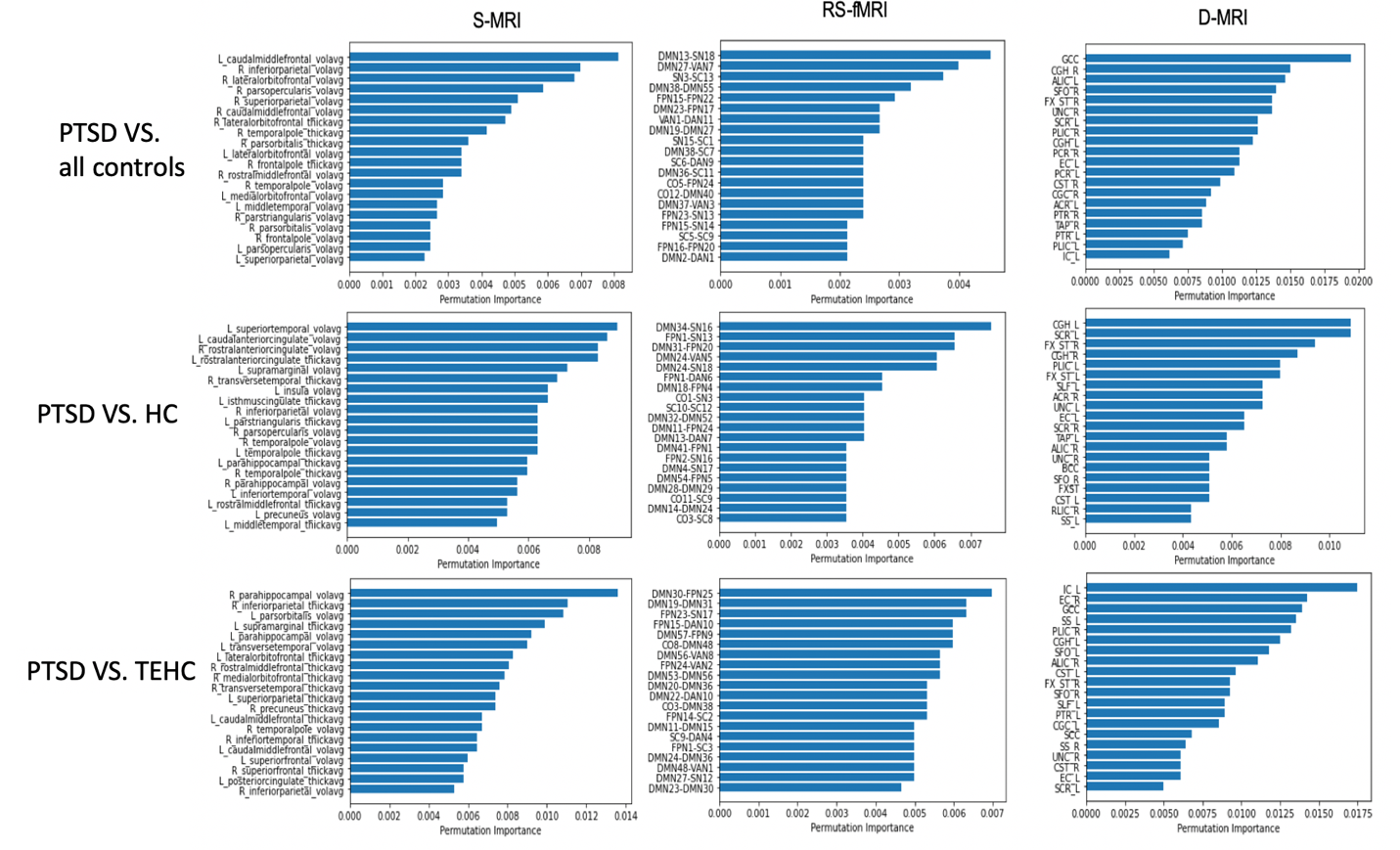
*Supplemental Figure 6: The feature importance using permutation importance method on a random forest classifier. Between PTSD and all controls (top row), PTSD and HC (middle row), and between PTSD and TEHC (bottom row), for s-MRI (first column), rs-fMRI (middle column), and d-MRI (third column)*

S-MRI

|  | CV AUC | Test AUC |
| --- | --- | --- |
| T1 RF | 0.59 | 0.6 |
| T1 SVM | 0.6 | 0.6 |
| T1 DVAE RF | 0.63 | 0.61 |
| T1 DVAE SVM | 0.63 | 0.62 |

RS-fMRI

|  | CV AUC | Test AUC |
| --- | --- | --- |
| RS RF | 0.57 | 0.58 |
| RS SVM | 0.59 | 0.59 |
| RS DVAE RF | 0.62 | 0.62 |
| RS DVAE SVM | 0.58 | 0.62 |

*Supplemental Figure 7: Compare classification performance using all features and DVAE-based latent variables in s-MRI (left) and rs-fMRI (right).*

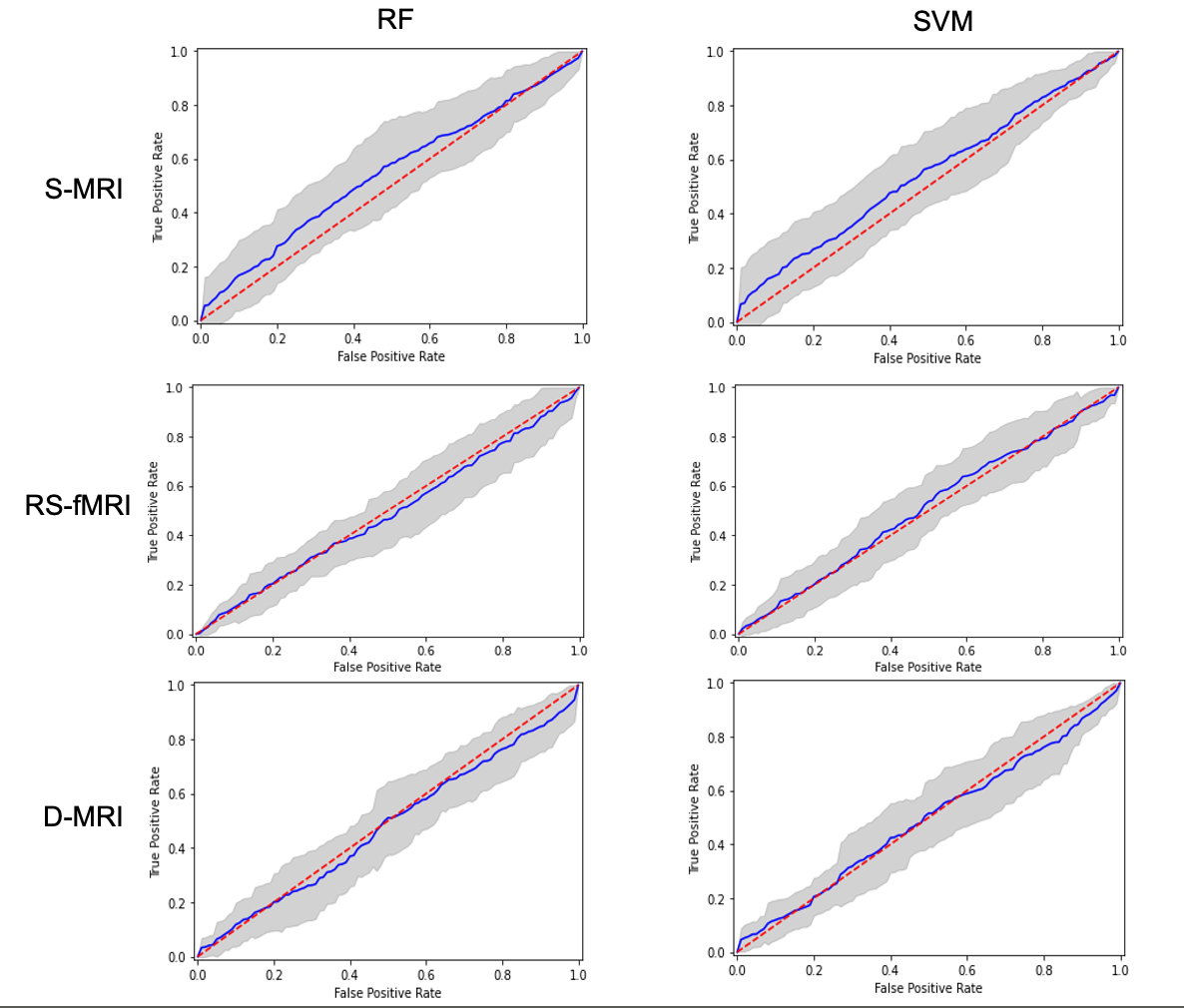

*Supplemental Figure 8: The AUC for leave one site out cross validation (LOSOCV) for s-MRI, rs-fMRI and d-MRI data using RF (left) and SVM (right)*

|  | Pooling | LOSOCV |
| --- | --- | --- |
| T1 | 0.65 | 0.56 |
| RS | 0.56 | 0.47 |
| DTI | 0.55 | 0.49 |
| T1 VAE | 0.63 | 0.63 |
| RS VAE | 0.62 | 0.62 |

*Supplemental Figure 9: The comparison of Leave one site out cross validation (LOSOCV) performance with aggregated pooling method across s-MRI (T1), rs-fMRI (RS), and d-MRI (DTI).*

Site

|  | Before | After |
| --- | --- | --- |
| s-MRI RF | 0.64 | 0.6 |
| s-MRI SVM | 0.63 | 0.52 |
| rs-fMRI RF | 0.54 | 0.54 |
| rs-fMRI SVM | 0.6 | 0.46 |
| d-MRI RF | 0.55 | 0.59 |
| d-MRI SVM | 0.56 | 0.52 |

*Supplemental Figure 10: Comparison of classification performance between PTSD and controls with controlling for sites using Combat, and without controlling for sites*

Age, Sex

|  | Before | After |
| --- | --- | --- |
| T1 RF | 0.64 | 0.61 |
| T1 SVM | 0.63 | 0.6 |
| RS RF | 0.54 | 0.54 |
| RS SVM | 0.6 | 0.6 |
| DTI RF | 0.55 | 0.55 |
| DTI SVM | 0.56 | 0.53 |

*Supplemental Figure 11: Comparison of classification performance between PTSD and controls without adding age and sex (Before), and with adding age and sex as features (After)*

*Supplemental Table 1: Demographic table for s-MRI by Sites:*

|  |  |  | **PTSD Diagnosis** | | **Sex** | | **Age** | | | |
| --- | --- | --- | --- | --- | --- | --- | --- | --- | --- | --- |
| **Site** | Full name | N | PTSD | Control | F | M | Min | Max | Mean | SD |
| **AMC** | Academic Medical Center | 75 | 38 | 37 | 35 | 40 | 22 | 59 | 39.99 | 9.9 |
| **ADNI-DOD** | ADNI-DOD | 194 | 80 | 114 | 1 | 193 | 61 | 85 | 69.04 | 4.77 |
| **Columbia** | Columbia U | 88 | 53 | 35 | 57 | 31 | 20 | 58 | 35.94 | 9.86 |
| **DeBellis** | Duke U - DeBellis | 118 | 30 | 88 | 64 | 54 | 6 | 16 | 10.31 | 2.59 |
| **Duke - Morey** | Duke U - Morey | 376 | 112 | 264 | 74 | 311 | 19 | 67 | 39.99 | 10 |
| **Ghent** | U of Ghent | 67 | 8 | 59 | 67 | 0 | 20 | 66 | 37.07 | 12.15 |
| **GTP** | Grady Trauma Project | 174 | 59 | 115 | 167 | 5 | 18 | 62 | 38.94 | 12.34 |
| **INTRuST** | INTRuST | 366 | 108 | 258 | 152 | 230 | 18 | 69 | 35.97 | 12.34 |
| **LUMC** | Leiden U. Medical Center | 52 | 22 | 30 | 45 | 7 | 12 | 20 | 15.25 | 1.81 |
| **McLean** | McLean Hospital-Kaufman | 52 | 39 | 13 | 52 | 0 | 19 | 62 | 37.52 | 12.32 |
| **McLean-Rosso** | McLean Hospital-Rosso | 113 | 22 | 91 | 61 | 53 | 20 | 51 | 34.06 | 8.99 |
| **Munster** | U Hospital Munster | 47 | 21 | 26 | 42 | 5 | 18 | 47 | 26.91 | 7.13 |
| **Utrecht** | U Medical Center Utrecht | 110 | 56 | 54 | 1 | 110 | 21 | 57 | 36.66 | 9.84 |
| **Irvine** | U of California Irvine | 28 | 13 | 15 | 0 | 28 | 25 | 48 | 33.54 | 7.15 |
| **CapeTown** | U of Cape Town | 51 | 7 | 44 | 67 | 0 | 20 | 48 | 29.33 | 6.86 |
| **UCAS** | U of Chinese Academy of Sciences | 70 | 34 | 36 | 39 | 31 | 37 | 61 | 49.56 | 6.82 |
| **Groningen** | U of Groningen | 40 | 40 | 0 | 40 | 0 | 23 | 58 | 38.17 | 9.69 |
| **Mannheim** | U of Mannheim | 49 | 49 | 0 | 49 | 0 | 20 | 63 | 35.86 | 11.64 |
| **Michigan** | U of Michigan | 69 | 26 | 43 | 11 | 58 | 19 | 63 | 31.09 | 10.96 |
| **UIUC** | U of Illinois at Chicago | 44 | 24 | 20 | 0 | 44 | 23 | 53 | 31.66 | 8.22 |
| **UMSL** | U of Missouri - St. Louis | 85 | 66 | 19 | 85 | 0 | 18 | 56 | 32.42 | 9.67 |
| **UNSW** | U of New South Wales | 167 | 49 | 118 | 102 | 65 | 18 | 69 | 40.23 | 12.63 |
| **SouthDakota** | U of South Dakota | 123 | 78 | 45 | 24 | 99 | 18 | 45 | 29.2 | 7.01 |
| **Toledo** | U of Toledo | 79 | 15 | 64 | 36 | 43 | 19 | 63 | 35.46 | 11.37 |
| **Uwash** | U of Washington | 246 | 54 | 192 | 130 | 125 | 8 | 20 | 13.92 | 3.06 |
| **Wisc-Madison** | U of Wisconsin - Madison | 58 | 19 | 39 | 4 | 54 | 22 | 48 | 30.64 | 6.39 |
| **Wisc-Milwaukee** | U of Wisconsin Milwaukee | 67 | 20 | 47 | 34 | 33 | 18 | 58 | 33.34 | 10.9 |
| **VA_AA** | VA Ann Arbor | 63 | 41 | 22 | 0 | 63 | 21 | 50 | 30.65 | 7.63 |
| **VAMinn** | VA Minneapolis | 243 | 94 | 149 | 13 | 230 | 22 | 60 | 32.71 | 7.93 |
| **Waco** | Waco | 66 | 41 | 25 | 10 | 56 | 25 | 60 | 40.85 | 11.15 |
| **WestHaven** | West Haven | 75 | 37 | 38 | 8 | 69 | 21 | 62 | 34.78 | 9.83 |
| **Yale** | Yale U | 72 | 23 | 49 | 11 | 60 | 20 | 52 | 30.06 | 7.86 |
|  | **Total** | **3527** | **1378** | **2149** | **1481** | **2097** |  |  |  |  |

*Supplemental Table 2: Demographic table for RS by Sites:*

|  |  |  | **PTSD Diagnosis** | |  | **Sex** | | **Age** | | | |
| --- | --- | --- | --- | --- | --- | --- | --- | --- | --- | --- | --- |
| **Site** | Full name | N | PTSD | Control | Control | M | F | Min | Max | Mean | SD |
|  |  |  |  |  | Type |  |  |  |  |  |  |
| **AMC** | Academic Medical Center | 70 | 34 | 36 | TEHC | 39 | 31 | 24 | 59 | 40.4 | 9.7 |
| **Beijing** | Beijing | 86 | 41 | 45 | TEHC | 37 | 49 | 22 | 66 | 49.1 | 10.2 |
| **CapeTown** | U of Cape Town /UCT | 108 | 5 | 103 | HC TEHC | 0 | 108 | 17 | 43 | 26.8 | 6.52 |
| **Columbia** | Columbia U | 43 | 8 | 35 | HC TEHC | 21 | 22 | 19 | 59 | 36.2 | 12.7 |
| **Duke** | Duke-Morey | 135 | 31 | 104 | TEHC | 102 | 33 | 21 | 67 | 38.7 | 10.6 |
| **Ghent** | U of Ghent | 65 | 8 | 57 | TEHC | 0 | 65 | 20.1 | 57.5 | 36.6 | 11.7 |
| **Groningen** | U of Groningen | 40 | 40 | 0 |  | 0 | 40 | 23 | 58 | 38.2 | 9.69 |
| **GTP** | Grady Trauma Project-Emory | 105 | 35 | 70 | TEHC | 0 | 105 | 18 | 62 | 38.5 | 11.4 |
| **LUMC** | Leiden U. Medical Center | 50 | 21 | 29 | HC | 6 | 44 | 12 | 20 | 15.2 | 1.8 |
| **Masaryk** | Masaryk U | 264 | 109 | 155 | TEHC | 104 | 160 | 15 | 94 | 51.7 | 18.8 |
| **McLean** | McLean Hospital-Kaufman/NTD | 64 | 42 | 22 | HC TEHC | 0 | 64 | 18 | 62 | 34.6 | 11.9 |
| **Michigan** | U of Michigan | 56 | 38 | 18 | HC TEHC | 56 | 0 | 22 | 50 | 31.3 | 7.8 |
| **Munster** | U Hospital Munster | 47 | 18 | 29 | HC | 9 | 20 | 18 | 51 | 25.8 | 6.81 |
| **Nanjing** | Nanjing Yixing | 129 | 45 | 84 | TEHC | 61 | 68 | 40 | 67 | 56.9 | 5.93 |
| **Stanford** | Stanford U | 180 | 99 | 81 | HC TEHC | 69 | 109 | 18 | 61 | 34.9 | 10.8 |
| **Toledo** | U of Toledo | 46 | 8 | 38 | TEHC | 26 | 20 | 19 | 63 | 34.4 | 11.6 |
| **Tours** | U of Tours | 39 | 9 | 30 | HC TEHC | 0 | 39 | 18 | 53 | 28.1 | 9.59 |
| **UMN** | U of Minnesota | 60 | 11 | 49 | TEHC | 55 | 5 | 25 | 61 | 42.3 | 9.65 |
| **Utrecht** | U Medical Center Utrecht | 104 | 53 | 51 | HC TEHC | 104 | 0 | 0 | 57 | 35.6 | 10.3 |
| **UWash** | U of Washington | 149 | 33 | 116 | HC TEHC | 75 | 74 | 8.08 | 17.3 | 12.8 | 2.65 |
| **VA_Minn** | VA Minneapolis | 247 | 90 | 157 | TEHC | 226 | 14 | 22 | 60 | 32.8 | 7.93 |
| **Vanderbilt** | Vanderbilt University Medical Center | 32 | 11 | 21 | HC TEHC | 26 | 6 | 23 | 40 | 31 | 4.72 |
| **Waco** | VA Waco | 22 | 9 | 13 | TEHC | 20 | 2 | 26 | 59 | 40.9 | 12.1 |
| **Westernontario** | Westernontario /Lawson | 158 | 106 | 52 | HC TEHC | 60 | 98 | 18 | 60 | 37.2 | 12.6 |
| **Wisc-Cisier** | U of Wisconsin - Madison | 98 | 83 | 15 | HC | 0 | 98 | 20 | 50 | 33.4 | 8.3 |
| **Wisc-Grupe** | U of Wisconsin - Madison-Grupe | 38 | 19 | 19 | TEHC | 34 | 4 | 22 | 46 | 31.1 | 6.53 |
| **Wisc-Milwaukee** | U of Wisconsin-Milwaukee/Wisc Larson | 67 | 18 | 49 | TEHC | 34 | 33 | 18.3 | 57.9 | 31.7 | 10.3 |
| **Total** | | **2502** | **1024** | **1478** |  | **1164** | **1311** |  |  |  |  |

*Supplemental Table 3: Demographic table for DTI by Sites:*

|  |  |  | **PTSD Diagnosis** | |  | **Sex** | | **Age** | | | |
| --- | --- | --- | --- | --- | --- | --- | --- | --- | --- | --- | --- |
| **Site** | **Site Full Name** | **N** | **PTSD** | **Control** | **Control type** | **Female** | **Male** | **Min** | **Max** | **Mean** | **SD** |
| **ADNIDOD** | ADNI-DOD | 135 | 69 | 66 | TEHC | 1 | 134 | 61 | 83 | 69.3 | 4.52 |
| **AMC** | Academic Medical Center | 70 | 34 | 36 | TEHC | 34 | 36 | 22 | 59 | 39.9 | 9.81 |
| **Beijing** | Beijing | 67 | 32 | 35 | TEHC | 35 | 32 | 37 | 61 | 49.4 | 6.88 |
| **Columbia** | Columbia U | 33 | 18 | 15 | TEHC | 33 | 0 | 22 | 53 | 33.58 | 8.18 |
| **Duke** | Duke U- Morey | 338 | 101 | 237 | HC/TEHC | 83 | 255 | 21 | 67 | 39.35 | 9.95 |
| **GTP** | Grady Trauma Project-Emory | 132 | 47 | 85 | HC/TEHC | 132 | 0 | 18 | 62 | 39.58 | 12.34 |
| **Groningen** | U of Groningen | 48 | 48 | NA | NA | 48 | 0 | 23 | 58 | 40.1 | 9.73 |
| **LUMC** | Leiden U. Medical Center | 40 | 20 | 20 | HC | 35 | 5 | 12 | 20 | 15.43 | 1.84 |
| **McLean** | McLean Hospital-Kaufman/NTD | 55 | 41 | 14 | HC/TEHC | 55 | 0 | 18 | 62 | 36.96 | 12.47 |
| **Munster** | U Hospital Munster | 4 | 2 | 2 | HC | 3 | 1 | 22 | 30 | 26 | 4.08 |
| **SouthDakota** | U of South Dakota | 91 | 55 | 36 | HC/TEHC | 9 | 82 | 22 | 45 | 31.79 | 6.13 |
| **Stellenbosch** | | 102 | 44 | 58 | TEHC | 63 | 39 | 21 | 77 | 44.52 | 13.32 |
| **CapeTown** | U of Cape Town /UCT | 57 | 8 | 49 | HC/TEHC | 57 | 0 | 20 | 48 | 29.4 | 6.75 |
| **Utrecht** | U Medical Center Utrecht | 94 | 46 | 48 | TEHC | 0 | 94 | 21 | 57 | 35.57 | 9.67 |
| **UWash** | U of Washington | 167 | 31 | 136 | TEHC | 86 | 81 | 8 | 19 | 14.52 | 2.84 |
| **VAMinn** | VA Minneapolis | 254 | 105 | 149 | HC/TEHC | 13 | 241 | 22 | 62 | 33.55 | 8.52 |
| **Waco** | Waco | 53 | 36 | 17 | HC | 7 | 46 | 25 | 60 | 39.58 | 10.83 |
| **Westernontario** | Westernontario /Lawson | 98 | 46 | 52 | TEHC | 46 | 52 | 18 | 59 | 34.7 | 12 |
| **Wisc_Grupe** | U of Wisconsin - Madison-Grupe | 48 | 16 | 32 | TEHC | 4 | 44 | 22 | 48 | 31 | 6.91 |
| **Yale** | Yale U | 67 | 37 | 30 | TEHC | 7 | 60 | 21 | 60 | 34.07 | 9.38 |
| **Total** |  | **1953** | **836** | **1117** |  | **751** | **1202** |  |  |  |  |

*Supplemental Table 4: inclusion and exclusion criteria for s-MRI, rs-fMRI and d-MRI data*

| Site | Cohort | Inclusion Criteria | Exclusion criteria |
| --- | --- | --- | --- |
| Academic Medical Center (AMC) | BOOSTER | PTSD patients had to fulfill the DSM-IV diagnostic criteria for PTSD, with a score of ≥ 45 on the clinician-administered PTSD scale (CAPS). Trauma-exposed controls: CAPS total score < 15. All: between 18-65 years of age, eligible for MRI. | PTSD patients were excluded if they met DSM-IV criteria for current psychotic disorder, substance-related disorder, severe personality disorder, severe major depressive disorder (MDD) (i.e., involving high suicidal risk and/or psychotic symptoms) or current suicidal risk. Trauma-exposed controls: no lifetime history of PTSD or MDD, or any current DSM-IV axis 1 disorder. All: no history of neurological disorders or any severe or chronic systemic disease or unstable medical condition, including endocrinological disorders. No use of psychotropic medications. Females: not pregnant or breastfeeding. |
| ADNI-DOD | ADNIDOD | PTSD: Subjects must be veterans of the Vietnam War, 50-90 years of age. Subjects who meet the SCID-I (for DSM-IV-TR) criteria for current/chronic PTSD (identified by records and verified by our telephone assessments). In addition to meeting DSM-IV-TR criteria for current/chronic PTSD, subjects must have a minimum current CAPS score of 50 as determined by telephone assessment. The PTSD symptoms contributing to the PTSD Diagnosis and current CAPS score must be related to a Vietnam War related trauma. Must live within 150 miles of the closest ADNI clinic in subject’s area. Controls: Subjects must be veterans of the Vietnam War, 50-90 years of age. Comparable in age, gender, and education with TBI and PTSD groups. May be receiving VA disability payments for something other than TBI or PTSD—or no disability at all. Must live within 150 miles of the closest ADNI clinic in subject’s area. | PTSD: Mild Cognitive Impairment/Dementia; Documented or self-report history of mild/moderate severe TBI; Any history of head trauma associated with injury onset cognitive complaints, or loss of consciousness for >5minutes. Controls: Mild Cognitive Impairment (MCI)/Dementia; Presence of PTSD by SCID-I for DSM-IV-TR criteria, or a CAPS score of >30 (Both current and/or a history of PTSD will be excluded); Documented or self-report history of mild/moderate severe TBI; Any history of head trauma associated with injury onset cognitive complaints, or loss of consciousness for >5 minutes; History of PTSD or current PTSD. Exclusionary criteria applied to TBI/PTSD will be applied to controls. All: MCI/dementia; History of psychosis or bipolar affective disorder; History of alcohol or substance abuse/dependence within the past 5 years (by DSM IV-TR criteria); MRI-related exclusions: aneurysm clips, metal implants that are determined to be unsafe for MRI, and/or claustrophobia; Contraindications for lumbar puncture, PET scan, or other procedures in this study; Seizure disorder or any systemic illness affecting brain function during the past 5 years; Clinical evidence of stroke; Having a history of relevant severe drug allergy or hypersensitivity; Subjects with current clinically significant unstable medical comorbidities, as indicated by history or physical exam, that pose a potential safety risk to the subject (Any major medical condition must be stable for at least 4 months prior to enrollment. These include but are not limited to clinically significant hepatic, renal, pulmonary, metabolic or endocrine disease, cancer, HIV infection and AIDS, as well as cardiovascular disease.). |
| Beijing | Wenchuan | age 18-65, personally experienced the earthquake | Major psychosis, drug or alcohol abuse |
| Columbia University | Columbia PE | Criterion A trauma. For patients, CAPS-4 diagnosis of PTSD, CAPS score of 50 or above. | For patients, psychosis, substance/alcohol dependence within 6 months or abuse within 2 months, use of psychotropic medication in past 4 weeks (6 weeks of fluoxetine), HAM-D-17 score greater than 24. For controls, current or past Axis I disorder or CAPS > 19. |
|  | Columbia Trauma (RS) | between the ages of 18 and 60. Experience of a traumatic event or events in childhood and/or adulthood; current DSM-V Criterion A for PTSD. Able to give consent, fluent in English | Prior or current Axis I psychiatric diagnosis of schizophrenia, psychotic disorder, bipolar disorder, dementia. Depression score of > 25 on the Hamilton Rating Scale for Depression (HAM-D-17-item); significant depression and /or depression related impairment that is judged to warrant pharmacotherapy or combined medication and psychotherapy.Individuals at risk for suicide based on history and current mental state. History of substance/alcohol dependence within the past six months, or abuse within past two months. Any psychotropic medications. Pregnancy, or plans to become pregnant during the period of the study. Paramagnetic metallic implants or devices contraindicating magnetic resonance imaging or any other non-removable paramagnetic metal in the body. Medical illness that could interfere with assessment of diagnosis, or biological measures (SCR, fMRI), including organic brain impairment from stroke, CNS tumor, or demyelinating disease; and renal, thyroid, hematologic or hepatic impairment. Any condition that would exclude MRI exam (e.g. pacemaker, paramagnetic metallic prosthesis, surgical clips, shrapnel, necessity for constant medicinal patch, some tattoos) |
| Duke University - DeBellis | DeBellis | Ages 6-14. | IQ<70; chronic medical illness; daily prescription medication; head injury with loss of consciousness; traumatic brain injury; neurological disorder; schizophrenia; anorexia nervosa; pervasive developmental disorder; obsessive compulsive disorder; bipolar I disorder or mania; birth weight under 5 lbs.; or severe prenatal (e.g. fetal alcohol and/or drug exposure) or perinatal complications (e.g. NICU stay); current or lifetime nicotine dependence/alcohol/substance use disorder; contraindications for safe MRI scan; and Axis I disorder or report of maltreatment that warranted CPS investigation in non-maltreated controls. |
| Duke University - Morey | CatGen | Age 18-65, fluent in English and capable of understanding consent, OEF/OIF Veteran. | Axis I disorders (except depression, GAD, PTSD, panic disorder, agoraphobia, other specific phobias, anxiety NOS), ferrous metal in the body, neurological disorders, history of TBI, color blindness, psychotic disorders, suicide attempts in past year, claustrophobia. |
|  | FearPTSD | Age 18-65, fluent in English and capable of understanding consent, OEF/OIF Veteran. | Axis I disorders (except depression, GAD, PTSD, panic disorder, agoraphobia, other specific phobias, anxiety NOS), ferrous metal in the body, neurological disorders, history of TBI, color blindness, psychotic disorders, suicide attempts in past year, claustrophobia. |
|  | MIRECC | Age 18-65, fluent in English and capable of understanding consent. | Axis I disorders (except depression, GAD, PTSD, panic disorder, agoraphobia, other specific phobias, anxiety NOS), ferrous metal in the body, neurological disorders, history of TBI, color blindness, psychotic disorders, suicide attempts in past year, claustrophobia. |
|  | Predator-1 | Age 18-65, fluent in English and capable of understanding consent, OEF/OIF Veteran. | Axis I disorders (except depression, GAD, PTSD, panic disorder, agoraphobia, other specific phobias, anxiety NOS), ferrous metal in the body, neurological disorders, history of TBI, color blindness, psychotic disorders, suicide attempts in past year, claustrophobia. |
|  | Predator-3 | Age 18-65, fluent in English and capable of understanding consent, OEF/OIF Veteran, free of implanted metal objects, unaffected by claustrophobia, and not be otherwise constrained from participating over the duration of the study in all related activities including scans and evaluations. | Significant neurological disorders, a history of learning disability, developmental delay, current substance abuse, a history of substance dependence, psychotic disorders, significant medical conditions, suicide attempt during the past year or are currently at high risk for suicide, neurological injury or disease (head trauma, seizures, strokes, prior neurosurgery, or if they are under the care of a neurologist or neurosurgeon), pregnant women, MRI contraindications. |
|  | Registry | Age 18-65, fluent in English and capable of understanding consent, OEF/OIF Veteran. | Ferrous metal in the body, claustrophobia. |
|  | SubBlast | Age 18-65, fluent in English and capable of understanding consent, OEF/OIF Veteran. | Axis I disorders (except depression, GAD, PTSD, panic disorder, agoraphobia, other specific phobias, anxiety NOS), ferrous metal in the body, neurological disorders, history of TBI, color blindness, psychotic disorders, suicide attempts in past year, claustrophobia. |
|  | TBIPTSD | Age 18-65, fluent in English and capable of understanding consent. | Axis I disorders (except depression, GAD, PTSD, panic disorder, agoraphobia, other specific phobias, anxiety NOS), ferrous metal in the body, neurological disorders, history of TBI, color blindness, psychotic disorders, suicide attempts in past year, claustrophobia. |
| University of Ghent | Ghent | For trauma group: experience(s) of physical, sexual, and/or emotional abuse occurring before 17 years of age as per SLESQ. For comparisons: no experience of childhood trauma and no experience of abuse-related trauma (e.g. emotional abuse, physical/sexual assault, etc.) later in life. For both: MRI compatibility (i.e., no pregnancy or metal implants), fluency in Dutch, normal or corrected-to-normal vision, female, and being 18-60 years of age. | History of severe head trauma or severe neurological condition. |
| Masaryk | Holocaust Survivors | Age 15-95, fluent in Czech/Slovak & capable of understanding consent, score in MMSE over 26 | Neurological disorders, claustrophobia, Psychotic disorders, ferrous metal in the body |
| Nanjing Yixing | Lost child | age 40-70, Chinese adults who had lost their only child | psychiatric disorders except PTSD (MDD, GAD); any history of or current brain injury or other major medical or neurological conditions, any MRI contraindication, left-handedness, unavailable data, and excessive head motion. We also excluded participants for whom the MRI scan was taken more than 120 months, or 10 years, after the child-loss event |
| Emory | GTP | African-American ethnicity; Female; English-speaking; Normal or corrected-to-normal vision. | Neurological disorder; history of major head injury; current psychosis; schizophrenia; bipolar disorder; psychotropic medication; illegal drug use (verified within 24 hours of scan with urine drug screen); pregnancy; typical safety criteria for MRI (metal implants etc.). |
| INTRuST | intrust | Patients: enrolled in individual INTRuST studies with a diagnosis of mTBI (initial Glasgow Coma Scale score of 13-15) or diagnosis of current psychological distress (PTSD, anxiety, or depression), or both. Healthy Controls: ages between 18 and 65. | Patients: (1) lifetime bipolar I disorder, lifetime psychotic disorders, lifetime dementia, delirium, alcohol or other substance dependence (within 30 days), (2) CNS disorders including aneurysm, anoxic events, brain tumor, encephalitis, Guillain Barre syndrome, Huntington's disease, hydrocephalus, uncontrolled diabetes, thyroid condition or blood pressure, multiple sclerosis, Parkinson's disease, seizure disorder, stroke, or subdural hematoma, (3) currently pregnant or lactating (due to effects of hormonal fluctuations on biological samples collected as part of the repository). (4) current medications that affect the brain function as determined by the study physician, (5) English as a second language after the age of 5, (6) history of a learning disability, and (7) weight of more than 300 pounds as this would preclude the subject from entering the scanner. Healthy Controls: screened by phone and by an in-person MINI (6.0.0) interview. (1) CNS disorders as described above, (2) medication exclusions, including more than one antihypertensive drug, psychotropic drugs within the last 90 days, herbal psychoactive substance use, or steroid use in the last 4 months, (3) currently pregnant or lactating, (4) history of mood, anxiety, psychotic, dementia, delirium, substance dependence in the past 12 months, (5) history of probable TBI as defined by the I-TBI. |
| Leiden University Medical Center | Episca | All participants met the following inclusion criteria: aged between 12 and 21, estimated full scale IQ (FIQ) ≥ 80 as measured by Dutch versions of the Wechsler Intelligence Scales for Children (WISC-III) or adults (WAIS), being right-handed, normal or corrected-to-normal vision, sufficient understanding of the Dutch language, no history of neurological impairments and no contraindications for MRI testing (e.g. braces, metal implants or possible pregnancy). | (1) primary DSM-IV diagnosis of ADHD, pervasive developmental disorders, Tourette’s syndrome, obsessive–compulsive disorder, bipolar disorder, and psychotic disorders, (2) current use of psychotropic medication other than stable use of SSRI’s, or amphetamine medication on the day of scanning, and (3) current substance abuse. |
| McLean Hospital - Kaufman | NTD | Female, age between 18 and 89, legal and mental competency, clinical diagnosis of PTSD with history of childhood abuse, or healthy controls. | Male; Under 18 or over 89; Legal or mental incompetence; Delirium secondary to medical illness; PTSD due to general medical or neurological illness; History of neurological conditions that may cause significant psychiatric symptomatology (e.g., dementia); Any contraindication to MR scans, including claustrophobia, pregnancy, metal implants, etc.; Current alcohol or substance dependence or abuse (within the last month); A history of schizophrenia or other psychotic disorder; History of head injury or loss of consciousness for longer than 5 min (including concussion). |
| McLean Hospital - Rosso | McLean Rosso | 1) 20-50 years of age; 2) right-handed; 3) DSM-IV diagnosis consistent with group assignment; 4) ability to provide written informed consent. | 1) Medical condition that would confound results; 2) history of seizures or head trauma with loss of consciousness; 3) exposure to psychotropic medications within 4 weeks of study (8 weeks for fluoxetine); 4) metal implants, claustrophobia or other MR exclusions; 5) positive urine toxicology or HCG status on scan day; 6) history of psychotic disorder, bipolar disorder, eating disorder, mental retardation, or pervasive developmental disorder; history of meeting full DSM-IV criteria for non-PTSD anxiety disorder. |
| Stanford University | BRAINS | Age 18-65, fluent in English & capable of understanding consent, OEF/OIF Veteran | History of psychotic, bipolar or substance dependence (within 3 months for patients and lifetime for controls), a history of a neurological disorder, greater than mild traumatic brain injury (i.e. >30 minutes loss of consciousness or >24 hour post-trauma amnesia), claustrophobia, and regular use of benzodiazepines, opiates, thyroid medications, or other CNS medication. Trauma-exposed healthy controls were required to have experienced a criterion A trauma, but not meet lifetime criteria for any Axis 1 psychiatric disorder, including PTSD. |
|  | CausCon | Patients are required to have chronic(>3 months)moderate to severe anxiety or depression, assessed dimensionally by a score on the PHQ9 scale(excluding the suicide question)>10 or a score on the GAD7scale >10. Both of these scales assess general symptoms of anxiety and depression, and these cutoffs have been shown to relate to moderate or greater severity of symptoms. Moreover, because these scales measure general anxiety and depression, they are sensitive to a wide range of DSM diagnoses, including GAD, MD and PTSD. Additionally, to ensure clinical significance, subjects will need to indicate that they would be interested in seeking treatment for these symptoms(i.e. that symptoms impair functioning). Other inclusion criteria are:(1)community dwelling adults ages 18-60 years old;(2)not currently in treatment;(3)free of metal or ferrous implant,(4)good English comprehension and non-impaired intellectual abilities to ensure understanding of task instructions;(5)no history of neurological disorders, brain surgery, electroconvulsive or radiation treatment, brain hemorrhage or tumor, stroke, epilepsy, hypo- or hyperthyroidism, and(6) no daily use of PRN benzodiazepines or opiates(max: 3x/wk), or daily thyroid medications, and no antidepressant, anticonvulsant or antipsychotic medications for > 2 wks(fluoxetine >6 wks). As-needed benzodiazepines or opiates cannot be used within 48 hours of assessments. Medication-free healthy subjects will likewise be split equally between those who have never been traumatized and those who have had a criterion A trauma. Controls must deny lifetime psychiatric diagnosis and treatment and have PHQ9 andGAD7≤4. Stratification of each group by trauma exposure will be re-assessed every 20 participants and wewill ensure that groups are matched on demographic variables. | 1. MRI counter-indications(e.g. shrapnel or other metal in/on the body that can not be removed, claustrophobia, etc.)2. Additional TMS counter-indications(seizure disorder, CNS active disorder, certain medications described below)3. Medication use that substantially reduces seizure threshold to TMS(olanzapine, chlorpromazine, lithium)and unwilling or unable medically(determined by patient and his/her physician)to safely withdraw, at least two weeks prior to TMS, from these medications4. Opiate medication, antihypertensive medication, or any medication that interferes with blood flow(interferes with fMRI recordings)5. Thyroid dysfunction not adequately controlled by medication6. History of neurological or cardiovascular disorders, brain surgery, radiation treatment, brain hemorrhage or tumor, stroke, or diabetes7. Diagnosis of substance dependence within the past 3 months(but not abuse)8. Refusal to abstain from illicit drug use for duration of the study9. Refusal to abstain from alcohol within 24 hours of scans10. Pregnancy in female participants11. Prior exposure to deep brain stimulation, rTMS, or tDCS(transcranial direct current stimulation)therapies12. Significant traumatic brain injury(loss of consciousness, post-injury amnesia, significant radiological/neurological findings, penetrating brain injury)13. Lifetime evidence of psychosis, mania, hypomania, or bipolar disorders on the SCID. |
| University Hospital Munster | Munster | All patients fulfilled the diagnostic criteria for PTSD as primary diagnosis according to the DSM-IV-TR (American Psychiatric Association, 2000), assessed by the German version of the Structured Clinical Interview for DSM-IV (SCID; Wittchen et al., 1997). Given the focus on InterPersonal Violence-PTSD (IPV-PTSD), the experience of a trauma related to IPV (e.g., rape, sexual or physical abuse) at least once was an inclusion criterion for the patient group. All participants had normal or corrected-to-normal vision and were right-handed as determined by the Edinburgh Handedness Inventory (Oldfield, 1971). | Controls: Lifetime PTSD. |
| University Medical Center Utrecht | BET | All: 18-60 years of age, eligible for MRI. PTSD: current PTSD diagnosis, with CAPS ≥ 45, military deployment >4 months. Trauma Controls: exposure to at least one traumatic event (according to DSM-IV A1 criterion), with CAPS < 15, no current psychiatric disorder, military deployment >4 months. Healthy Controls: no current psychiatric disorder according to DSM-IV. | All: history of neurological disorders, any severe or chronic disorder; alcohol or drug abuse and/or dependence during course of the study. |
| University of California Irvine | UCI | General: 1) male or female between 18-65 years of age, 2) IQ level > 75, and 3) able to communicate effectively with the investigator and study coordinator and the ability to provide informed consent. PTSD Patients: 1) OEF/OIF/OND veterans with DSM-IV criteria for chronic PTSD on the Clinician Administered PTSD Scale (CAPS-4) and 2) at least moderate PTSD by having a total CAPS score of > 50. Eligible persons will be allowed to have other symptoms that are commonly comorbid with PTSD (e.g., anxiety, mild to moderate depression, somatic symptoms), but these will not be inclusion or exclusion criteria. | General: 1) over scanner weight and size limits or insufficient visual acuity to view visual stimuli in the scanner, 2) contra-indications for MRI scans: pacemaker, metal implants, tattoos on face/neck, positive urine pregnancy test, claustrophobia etc., 3) current or past history of major medical illness that may affect brain function, 4) Other DSM-IV axis 1 disorders (history of substance abuse or dependence in full remission is acceptable), 5) positive urine drug screen, 6) history of drug dependence in the past 3 years, or current substance abuse, or 7) any active medical condition that in the investigator’s opinion will interfere with the study’s objectives. Controls: a first-degree relative (parent, sibling, child) with an Axis 1 anxiety diagnosis. PTSD Patients: 1) psychosis, 2) substance dependence within the past 6 months, 3) thyroid disease, 4) decisional incapacity (e.g., dementia), 5) centrally acting medications that potentially have an effect on biological expression (e.g., beta-blockers, opiates, and >10mg equivalent of diazepam/day), 6) pain levels requiring opiate medications, 7) known exposure to chemicals or physical trauma that cause neuropsychiatric sequelae, 8) severe depression (Beck Depression Inventory-II3 score >30) since this may bias accurate PTSD diagnosis and biological measures, 9) a diagnosed sleep breathing disorder, 10) past chronic PTSD prior to military service. |
| University of Cape Town | Drakenstein Child Health Study | Women over the age of 18 years, who were between 20 and 28 weeks pregnant at the time of initial inclusion in the study, who presented to one of two health care clinics for antenatal care (TC Newman and Mbekweni clinics), and had no intention of moving out of the area within the following year, and were able to give written consent | 1) loss of consciousness longer than 30 minutes, 2) inability to  speak English, 3) current/lifetime alcohol and/or substance dependence or abuse, 4)  psychopathology other than PTSD and/or MDD, 5) traumatic brain injury, 6) standard MRI  exclusion criteria, such as claustrophobia and presence of ferromagnetic objects in the  participant’s body |
| University of Chinese Academy of Sciences | UCAS | (1) household was used as the basic sample unit, and only one member within each household was randomly selected for participation; (2) eligible participants included those who were at least 16 years old, and experienced the disaster personally; (3) among the eligible participants, the household member whose birthday was closest to the date of investigation was first selected for participation, and if the individual was unavailable, the household member whose birthday was the next closest was selected; this procedure continued until a participant was identified. | Individuals with mental disability and any major psychosis (e.g., schizophrenia and organic mental disorders) were excluded. |
| University of Groningen | DISPO (RS) | Age 20-60, ALL FEMALES with trauma exposure before age 21; Diagnostic interviews by Clinician with CAPS, SCID-1, SCID-2, and SCID-D; all subjects have PTSD diagnosis (according to CAPS), fluent in German, | Psychotic disorders, dissociative identity disorder, current alcohol dependency, current substance intoxication, ferrous metal in the body, neurological disorders, history of TBI, claustrophobia |
| University of Mannheim | Mannheim | PTSD: Women aged 18-65 years, PTSD after childhood sexual or physical abuse before the age of 18 years, Sexual or physical assault must be the index trauma, At least 3 criteria of BPD (including criterion 6: affective instability; IPDE), Commitment and possibility to attend weekly therapy sessions for one year, No planned absence for more than 4 weeks in this period. Trauma Controls: Women aged 18-65 years, Childhood sexual or physical abuse before the age of 18 years. Healthy Controls: Women aged 18-65 years. | PTSD: general exclusion criteria were traumatic brain injuries, current and lifetime schizophrenia or bipolar-I disorder, mental retardation, severe psychopathology or somatic illness that needs to be treated immediately in another setting (e.g., BMI<16), medical conditions making exposure-based treatment impossible, a suicide attempt within the last two months, and substance dependency with no abstinence within two months prior to the study. Trauma Controls and Healthy Controls: any current or previous mental disorder, any psychotherapeutic experience or any intake of psychotropic medication during the lifetime or presently. For the current fMRI study, further exclusion criteria were metal implants, pregnancy, left-handedness, and claustrophobia. |
| University of Michigan | CCMB08 | All: 18-55 years of age, combat veterans and civilians, eligible for MRI. PTSD: current PTSD diagnosis, with CAPS ≥ 50. Trauma Controls: exposure to at least one traumatic event (according to DSM-IV A1 criterion), with CAPS < 15. Community Controls: CAPS < 15. | All: history of neurological disorders, any severe or chronic disorder; alcohol or drug abuse and/or dependence during course of the study. |
|  | R24 | All: 18-55 years of age, combat veterans and civilians, eligible for MRI. PTSD: current PTSD diagnosis, with CAPS ≥ 50. Trauma Controls: exposure to at least one traumatic event (according to DSM-IV A1 criterion), with CAPS < 15. Community Controls: CAPS < 15. | All: history of neurological disorders, any severe or chronic disorder; alcohol or drug abuse and/or dependence during course of the study. |
|  | Mindfulness (RS) | age 18-65, fluent in English & capable of understanding consent, OEF/OIF Veteran | Axis I disorders (except Depression, GAD, PTSD, Panic Disorder, Agoraphobia, Other Specific Phobias, Anxiety NOS), neurological disorders, Current or history of Psychotic disorders, Suicide attempts in past year, ferrous metal in the body,, claustrophobia, or other contraindication for MRI, |
| University of Tours | COPTSD (RS) | Age 18-65, fluent in French, scanning time at 6 months post exposure to sexual assault. | History of head injury, substance use, claustrophobia, current use of a psychotropic medication for more than 21 days, medical disorders affecting brain function (e.g., epi- lepsy, tumour), and MRI contraindications. |
| University of Illinois at Chicago | Phan | All: 18-55 years of age, discharge from active military service (post OEF/OIF deployment), ability to read and speak English. PTSD: current PTSD related to combat trauma, CAPS ≥ 40 with greater than 1 month duration of symptoms, CES score ≥ 17. Combat Controls: CAPS < 20, CES score ≥ 17. | All: life history of bipolar disorder, schizophrenia, mental retardation, passive developmental disorder; severe depressive symptoms as indicated by HAM-D score of ≥ 25; current alcohol/drug dependence within the past 6 months; current suicidal/homicidal ideation; ongoing psychotherapy treatment; left-handedness. PTSD: presence of clinically significant medical condition or taking a medication which interferes with metabolism of paroxetine; history of hypersensitivity to paroxetine/SSRI; prior failure of response to paroxetine/SSRI for PTSD; history of PTSD or partial/subthreshold PTSD related to a prior deployment or a prior trauma. |
| University of Minnesota (UMN) | Mars2 | Age 18-65, history of combat-related trauma meeting DSM-5, criterion-A stressors | Psychiatric health: Current or past history of any psychotic disorder, history of any psychotic disorder, bipolar disorder, delirium, dementia, amnestic disorder, or mental retardation; comorbid depression if accompanied by current, significant suicide risk; substance use disorder presently or for the six months preceding testing. Medical health and pregnancy status: Current or past medical illnesses which in the investigator’s opinion may confound study results, or place the participant at risk; Females who are, or may be, pregnant. Medication: Current use of any medication that alters central nervous system function including antidepressants, benzodiazipines, anti-psychotics, mood-stabilizers, anti-parkinsonian agents, anti-convulsants, sleep medications, pain medications, and anti-hypertensives. MRI Safety: Ferrous metal in the body, other MRI contraindication |
| University of Missouri – St. Louis | Bruce | Patients met DSM-IV criteria for PTSD that resulted from interpersonal trauma and had a CAPS score of 45 or higher. For controls, no Criterion A event or current diagnosis of mood or anxiety disorder. | Psychosis, bipolar disorder, current psychotropic medication use, current diagnosis of substance of alcohol abuse, history of head trauma or neurological disorders, MRI contraindications. |
| University of New South Wales | CCRE | DSM-IV (CAPS). | Moderate/severe TBI; Substance dependence; History of psychosis; History of neurological disorder. |
| University of South Dakota | PTSD | OIF/OEF/OND (Operation New Dawn) veterans. | Exclusion criteria were (a) current or previous seizure history; b) current crisis-related issues such as serious self-injurious behavior, psychosis, or substance dependence (excluding alcohol dependence); (c) report of traumatic brain injury using the Traumatic Brain Injury Checklist; and (d) contraindications to fMRI (metal objects in body, claustrophobia). |
|  | SAP | Participants were undergraduate students who were identified as an adult child of an alcoholic parent (ACoA), based on the Children of Alcoholics Screening Test (CAST, Jones, 1983). A score of 6 or above on the CAST indicated the participant was more than likely the child of an alcoholic parent and raised by this parent. | Participants were excluded for current or previous seizure history, contraindications to MRI, or if they exhibited possible psychotic or other psychological symptoms that would make inclusion in the study potentially hazardous to them. |
| University of Toledo | MVA | Survivors of a Motor Vehicle Accident (MVA) who are transported to the University of Toledo Emergency department, or to a ProMedica emergency medicine department. | Pregnancy; under the influence of alcohol or drugs at the time of MVA; major injuries, moderate to severe traumatic brain injury; major medical illnesses; conditions affecting ability to undergo MRI scans. |
|  | ONG | Ohio National Guard and Reserve soldiers, deployment in OEF or OIF; 18-50 years of age, meeting Ohio National Guard Study characteristics, and able to give informed consent. | a) life history of a neurologic condition, psychosis or bipolar disorder, b) active substance dependence, c) organic mental syndrome or pervasive developmental disorder; d) presence of ferrous metal in the body (e.g., aneurysm clips, shrapnel); or e) no psychotropic medication is preferred while use of antipsychotics will be excluded. |
| University of Washington | FEAR | Aged 8-17. | Psychiatric medication use (excepting stimulant meds for ADHD), braces, claustrophobia, active substance dependence, pervasive developmental disorder, non-English speaking, active safety concerns. |
|  | NARSAD | Aged 8-19. | Psychiatric medication use (excepting stimulant meds for ADHD), braces, claustrophobia, active substance dependence, pervasive developmental disorder, non-English speaking, active safety concerns. |
|  | SAS | Aged 13-20. | Psychiatric medication use (excepting stimulant meds for ADHD), braces, claustrophobia, active substance dependence, pervasive developmental disorder, non-English speaking, active safety concerns. |
|  | MT (RS) | 8-20 years old English speaking | Psychiatric medication use (excepting stimulant meds for ADHD, which were discontinued for the scan). MRI contra-indications including braces, or other metal in the body or claustrophobia. Active substance dependence, pervasive developmental disorder, active safety concerns. |
| University of Wisconsin - Madison | UWMadison | Age range of 18-50; Capable of giving informed consent; Fluent in English; Exposure to one or more life-threatening war zone trauma events per the Combat Experiences Scale and documented by DD-214, Combat Action Ribbon (Marines), Combat Infantry Badge (Army), or other clear evidence of war zone trauma exposure in Iraq or Afghanistan since 2001; Pharmacological or psychotherapeutic treatment stable for at least 8 weeks prior to beginning of study, with no intent to begin a new course of treatment during the study period. | Weight of 352 pounds or over (due to constraints of MRI scanner); Women of childbearing potential with positive pregnancy test, looking to conceive during the research timeline, or who are breastfeeding; Metallic implants such as prostheses or aneurysm clip, or electronic implants such as cardiac pacemakers; Neurological or serious medical condition that may contraindicate MRI or that may overlap with physiological substrates of psychiatric conditions; History of seizures or seizure disorder; Moderate or severe traumatic brain injury (over 20 minutes unconscious); Current active substance dependence or dependence within 3 months (other than nicotine); Meets DSM-IV criteria for bipolar disorder, schizophrenia, schizoaffective disorder, psychotic disorder NOS, delirium, or any DSM-IV cognitive disorder; Substance dependence disorder within 3 months or any current substance dependence; Severe psychiatric instability or severe situational life crises, including evidence of being actively suicidal or homicidal, or any behavior that poses an immediate danger to patient or others; Participants with extensive experience in yoga or meditation; Current use of benzodiazepines and beta-blockers. |
| University of Wisconsin - Madison-Grupe | Veterans' Wellness | Age range of 18-50. Capable of giving informed consent Fluent in English Exposure to one or more life-threatening war zone trauma events per the Combat Experiences Scale and documented by DD-214, Combat Action Ribbon (Marines), Combat Infantry Badge (Army), or other clear evidence of war zone trauma exposure in Iraq or Afghanistan since 2001 Pharmacological or psychotherapeutic treatment stable for at least 8 weeks prior to beginning of study, with no intent to begin a new course of treatment during the study period | Medical Weight of 352 pounds or over (due to constraints of MRI scanner) Women of childbearing potential with positive pregnancy test, looking to conceive during the research timeline, or who are breastfeeding Metallic implants such as prostheses or aneurysm clip, or electronic implants such as cardiac pacemakers Neurological or serious medical condition that may contraindicate MRI or that may overlap with physiological substrates of psychiatric conditions History of seizures or seizure disorder Moderate or severe traumatic brain injury (over 20 minutes unconscious)   Psychiatric/Behavioral Current active substance dependence or dependence within 3 months (other than nicotine) Meets DSM-IV criteria for bipolar disorder, schizophrenia, schizoaffective disorder, psychotic disorder NOS, delirium, or any DSM-IV cognitive disorder. Substance dependence disorder within 3 months or any current substance dependence Severe psychiatric instability or severe situational life crises, including evidence of being actively suicidal or homicidal, or any behavior that poses an immediate danger to patient or others. Participants with extensive experience in yoga or meditation  Medications/Therapies Current use of benzodiazepines and beta-blockers |
| University of Wisconsin - Milwaukee | WiscLarson | PTSD criterion A met, age 18-60, GCS>= 13 (mild TBI criteria), Rothbaum 3 or higher or item 2 rated 3 or higher, English speaking (either native or bilingual proficiency), able to schedule within 30 days of brain injury. | Still in high school, re-admitted to hospital for current brain injury, live too far away to travel for study, police hold, incarcerated, intentional self-inflicted injury, known perpetrator, moderate to severe cognitive impairment, loss of consciousness > 30 minutes, pregnant, clear evidence of substance abuse, anti-psychotic or anti-seizure medication, indication of psychotic disorder or manic symptoms, MRI contraindications, history of seizures or other neurological conditions, severe hearing or vision problems. |
| University of Wisconsin-Madison /Wisc_Cisler | DOP | Age 21-50, fluent in English, experience of interpersonal violence | Psychotic symptoms, past psychotic disorders, stable on medications < 4 weeks, cognitive impairment, current substance or alcohol use disorder |
|  | EMOREG | Age 21-50, fluent in English, experience of interpersonal violence (for PTSD group) | Psychotic symptoms, past psychotic disorders, stable on medications < 4 weeks, cognitive impairment, current substance or alcohol use disorder |
|  | PAL | Age 21-50, fluent in English, experience of interpersonal violence (for PTSD group) | Psychotic symptoms, past psychotic disorders, stable on medications < 4 weeks, cognitive impairment, current substance or alcohol use disorder |
| Vanderbilt | Vanderbilt PTSD Study | age 18-50, English speaking, OEF/OIF/OND Veteran | Individuals in the TEHC and healthy control (HC) groups will not have current nor past PTSD. Furthermore, the TEHC and HC groups will not have symptoms of hypervigilance. Individuals in the HC group will not have exposure to a traumatic event. In all groups, individuals will be excluded for: 1) Current use of psychoactive medications (within past 6 weeks) or current psychological therapy (> 1 month) 2) Current substance use disorder (must be in remission at least 6 months). Subjects will be required to screen negative on a urine drug screen (Triage Drugs of Abuse Panel, Biosite Diagnostics, San Diego, CA) and alcohol breath screen (Intoximeters, Inc, St. Louis, Missouri) on the MRI study day; 3) A psychotic disorder or bipolar disorder, determined using the Structured Clinical Interview for DSM-IV (First, Spitzer, Gibbon, & JBW, 2002) Lifetime.  4) Current or past traumatic brain injury (TBI) 5) Significant medical illness (e.g., cancer, HIV), or neurological illness (e.g., stroke, brain tumor, multiple sclerosis, epilepsy)  6) MRI safety risk (e.g., metal in body). |
| VA Ann Arbor | Mindfulness | Inclusion criteria were long‐term (>10 years) PTSD as assessed by Clinician Administered PTSD Scale, or PTSD in partial remission. | Exclusion criteria included diagnoses of psychosis (e.g. schizophrenia, bipolar, and schizoaffective disorders) and current substance dependence, or active suicidal intent, as assessed using the Mini International Neuropsychiatric Interview (MINI). |
| VA Minneapolis VA (VAminn) | SATURN | age: 18-60, OEF/OIF, deployed | Moderate/severe TBI, non-TBI neurological conditions, current psychotic symptoms, substance abuse/dependence other than alcohol, unstable med conditions, sig risk of suicide/homicide |
|  | DEFEND | age: 18-60, OEF/OIF, deployed, positive screen on VA TBI Clinical Reminder | mod/sev TBI, non-TBI neurological conditions, current psychotic symptoms, substance abuse/dependence other than alcohol, unstable med conditions, sig risk of suicide/homicide |
| VA Waco | MAVERIX | Veteran, age 18-60, agreement to donate saliva | Serious general medical condition that would risk the subject being able to complete MRI (active seizure disorder, dementia, active back or muscle spasms), MRI safety screen positive (metal) or history of penetrating head or eye wound without subsequent radiological evidence that the wound is metal-free. Subjects that are (were) welders or subjects that have had metal surgically removed from their eyes will not be allowed to participate without subsequent radiological evidence that the wound is metal-free, MRI quality problems (tremors, significant claustrophobia, teeth braces) |
|  | ROBI | Participants will be Veterans with a diagnosis of TBI, recruited on a volunteer basis from the Central Texas VA. Inclusion criteria are age of 18-60 years and a clinical diagnosis of TBI in the VA medical record. | Exclusion criteria will include: an absence of qEEG parameters more than 2 standard deviations from the population mean of healthy age-matched historical controls saved in a commercial normative database (Neuroguide, Largo, FL); a positive screen on the MINI International Neuropsychiatric Interview: diagnosis of schizophrenia, schizoaffective disorder, bipolar disorder type I, severe substance use disorder, a high risk of suicide; and an inability to provide informed consent. |
|  | TEMI | Male and female Veterans enrolled in a CTVHCS PTSD treatment program who are 18-60 years old. | (1) pregnancy; (2) exposure to metal in the eyes; (3) shrapnel or other metal embedded in the body; (4) ferromagnetic surgical implants; (5) mechanical implants (e.g., pacemakers); (6) electrical implants (e.g., cochlear implants); (7) non-removable metallic devices (e.g. stables, neck braces, or artificial limbs) (8) tattoos not done professionally; (9) non-removable body piercings; (10) current psychosis including Axis I psychotic disorder, bipolar disorder, or schizophrenia; (11) dementia or another severe cognitive disorder; (12) prior exposure to an rTMS or dTMS; (13) seizure disorder; (14) positive screen for suicidal intent, plan, or behavior within the past 6 months; (15) a TMS motor threshold of 70% or greater of the machine’s maximum output. |
| West Haven | West Haven | Combat-exposed veterans with PTSD and age-matched male combat-exposed healthy controls (combat controls). All participants had been deployed on one or more tours to Iraq and/or Afghanistan and reported exposure to combat-related experiences. All participants were 18 to 50 years of age. | Participants were excluded based on moderate and severe TBI, neurological disorder, and MRI contraindications. Participants with PTSD were also excluded on the basis of a diagnosis of current drug/alcohol abuse, recent change in antidepressant medications (stable dose for 4 weeks required). |
| Western Ontario | Lawson (RS) | Primary diagnosis PTSD | Psychotic disorder, bipolar disorder, traumatic brain injury, narcotic use, active substance use disorder within 3 months of study entry |
| Yale | Yale | Participants ranged in age from 21 to 60 and had been deployed on one or more combat tours. | Individuals were excluded from the study if they met any of the following criteria: a diagnosis of bipolar disorder or psychotic disorder, as assessed by the SCID-IV (First, Spitzer, Gibbon, & Williams, 2002); current benzodiazepine use; a history of ADHD, learning disorder, moderate or severe traumatic brain injury (TBI), brain tumor, epilepsy, or a neurological disorder; current inpatient status; or an MRI contraindication. |

*Supplemental Table 5: List of features from structural MRI.*

*abbreviations, L means Left, R means Right, thickavg: averaged cortical thickness in this ROI, volavg: averaged volume in each ROI*

| ROI | Structural ROI | Full name | Lobe |
| --- | --- | --- | --- |
| 1 | Caudalanteriorcingulate (L+R) | Caudal anterior cingulate | Cingulate /Frontal |
| 2 | Caudalmiddlefrontal (L+R) | Caudal middle frontal | Frontal |
| 3 | Inferiorparietal (L+R) | Inferior parietal | Parietal |
| 4 | Inferiortemporal (L+R) | Inferior temporal | Temporal |
| 5 | Isthmuscingulate (L+R) | isthmus cingulate | Cingulate /Parietal |
| 6 | Lateralorbitofrontal (L+R) | lateral orbitofrontal | Frontal |
| 7 | Medialorbitofrontal (L+R) | medial orbitofrontal | Frontal |
| 8 | Middletemporal (L+R) | middle temporal | Temporal |
| 9 | Parahippocampal (L+R) | parahippocampal | Temporal/Limbic |
| 10 | Parsopercularis (L+R) | parsopercularis | Frontal |
| 11 | Parsorbitalis (L+R) | parsorbitalis | Frontal |
| 12 | Parstriangularis (L+R) | parstriangularis | Frontal |
| 13 | Posteriorcingulate (L+R) | posterior cingulate | Cingulate /Parietal |
| 14 | Precuneus (L+R) | precuneus | Parietal |
| 15 | Rostralanteriorcingulate (L+R) | rostral anterior cingulate | Cingulate /Frontal |
| 16 | Rostralmiddlefrontal (L+R) | rostral middle frontal | Frontal |
| 17 | Superiorfrontal (L+R) | superior frontal | Frontal |
| 18 | Superiorparietal (L+R) | superior parietal | Parietal |
| 19 | Superiortemporal (L+R) | superior temporal | Temporal |
| 20 | Supramarginal (L+R) | supramarginal | Parietal |
| 21 | Frontalpole (L+R) | Frontal pole | Frontal |
| 22 | Temporalpole (L+R) | Temporal pole | Temporal |
| 23 | Transversetemporal (L+R) | Transverse temporal | Temporal |
| 24 | Insula (L+R) | Insula |  |

*Supplemental Table 6: Rs-fMRI ROI definition by Power atlas, network assignment and abbreviations:*

|  |  |  | MNI space | | |  |  |
| --- | --- | --- | --- | --- | --- | --- | --- |
| Suggested System | Network Label | Power ROI | X | Y | Z | Harvard-Oxford Cortical Structural Atlas | Harvard-Oxford Subcortical Structural Atlas |
| Cingulo-opercular Task Control | CO1 | 47 | -3 | 2 | 53 | 82% Juxtapositional Lobule Cortex (formerly Supplementary Motor Cortex) |  |
| Cingulo-opercular Task Control | CO10 | 56 | 49 | 8 | -1 | 75% Central Opercular Cortex |  |
| Cingulo-opercular Task Control | CO11 | 57 | # | 3 | 4 | 10% Insular Cortex |  |
| Cingulo-opercular Task Control | CO12 | 58 | # | 8 | -2 | 35% Central Opercular Cortex 30% Precentral Gyrus 6% Inferior Frontal Gyrus, pars opercularis 5% Frontal Operculum Cortex |  |
| Cingulo-opercular Task Control | CO13 | 59 | -5 | 18 | 34 | 49% Cingulate Gyrus, anterior division 30% Paracingulate Gyrus |  |
| Cingulo-opercular Task Control | CO14 | 60 | 36 | 10 | 1 | 42% Insular Cortex |  |
| Cingulo-opercular Task Control | CO2 | 48 | 54 | # | 34 | 16% Supramarginal Gyrus, anterior division 10% Parietal Operculum Cortex 5% Supramarginal Gyrus, posterior division |  |
| Cingulo-opercular Task Control | CO3 | 49 | 19 | -8 | 64 | 19% Precentral Gyrus 15% Superior Frontal Gyrus |  |
| Cingulo-opercular Task Control | CO4 | 50 | # | -5 | 71 | 42% Superior Frontal Gyrus 9% Precentral Gyrus |  |
| Cingulo-opercular Task Control | CO5 | 51 | # | -2 | 42 | 28% Juxtapositional Lobule Cortex (formerly Supplementary Motor Cortex) 14% Cingulate Gyrus, anterior division |  |
| Cingulo-opercular Task Control | CO6 | 52 | 37 | 1 | -4 | 6% Insular Cortex | 10% Right Putamen |
| Cingulo-opercular Task Control | CO7 | 53 | 13 | -1 | 70 | 27% Superior Frontal Gyrus 14% Juxtapositional Lobule Cortex (formerly Supplementary Motor Cortex) 7% Precentral Gyrus |  |
| Cingulo-opercular Task Control | CO8 | 54 | 7 | 8 | 51 | 80% Juxtapositional Lobule Cortex (formerly Supplementary Motor Cortex) 8% Paracingulate Gyrus 6% Cingulate Gyrus, anterior division |  |
| Cingulo-opercular Task Control | CO9 | 55 | # | 0 | 9 | 71% Central Opercular Cortex 12% Insular Cortex |  |
| Default mode | DMN1 | 74 | # | # | 26 | 53% Lateral Occipital Cortex, superior division |  |
| Default mode | DMN10 | 83 | # | # | # | 74% Middle Temporal Gyrus, posterior division |  |
| Default mode | DMN11 | 86 | # | # | 35 | 55% Lateral Occipital Cortex, superior division 7% Angular Gyrus |  |
| Default mode | DMN12 | 87 | # | # | 44 | 60% Lateral Occipital Cortex, superior division |  |
| Default mode | DMN13 | 88 | -7 | # | 27 | 35% Cingulate Gyrus, posterior division 34% Precuneous Cortex |  |
| Default mode | DMN14 | 89 | 6 | # | 35 | 69% Precuneous Cortex |  |
| Default mode | DMN15 | 90 | # | # | 16 | 51% Precuneous Cortex 6% Supracalcarine Cortex |  |
| Default mode | DMN16 | 91 | -3 | # | 13 | 62% Cingulate Gyrus, posterior division 19% Precuneous Cortex |  |
| Default mode | DMN17 | 92 | 8 | # | 31 | 42% Cingulate Gyrus, posterior division 20% Precuneous Cortex |  |
| Default mode | DMN18 | 93 | 15 | # | 26 | 47% Precuneous Cortex 24% Supracalcarine Cortex 10% Cuneal Cortex |  |
| Default mode | DMN19 | 94 | -2 | # | 44 | 68% Cingulate Gyrus, posterior division 27% Precuneous Cortex |  |
| Default mode | DMN2 | 75 | 6 | 67 | -4 | 90% Frontal Pole |  |
| Default mode | DMN20 | 95 | 11 | # | 17 | 51% Precuneous Cortex 7% Supracalcarine Cortex |  |
| Default mode | DMN21 | 96 | 52 | # | 36 | 42% Lateral Occipital Cortex, superior division 15% Angular Gyrus |  |
| Default mode | DMN22 | 97 | 23 | 33 | 48 | 45% Superior Frontal Gyrus 6% Middle Frontal Gyrus |  |
| Default mode | DMN23 | 98 | # | 39 | 52 | 66% Superior Frontal Gyrus 6% Frontal Pole |  |
| Default mode | DMN24 | 99 | # | 29 | 53 | 30% Superior Frontal Gyrus |  |
| Default mode | DMN25 | 100 | # | 20 | 51 | 35% Middle Frontal Gyrus |  |
| Default mode | DMN26 | 101 | 22 | 39 | 39 | 36% Superior Frontal Gyrus 16% Frontal Pole 11% Middle Frontal Gyrus |  |
| Default mode | DMN27 | 102 | 13 | 55 | 38 | 74% Frontal Pole |  |
| Default mode | DMN28 | 103 | # | 55 | 39 | 77% Frontal Pole |  |
| Default mode | DMN29 | 104 | # | 45 | 39 | 40% Frontal Pole 22% Superior Frontal Gyrus 5% Middle Frontal Gyrus |  |
| Default mode | DMN3 | 76 | 8 | 48 | # | 44% Paracingulate Gyrus 26% Frontal Medial Cortex |  |
| Default mode | DMN30 | 105 | 6 | 54 | 16 | 75% Paracingulate Gyrus 11% Superior Frontal Gyrus |  |
| Default mode | DMN31 | 106 | 6 | 64 | 22 | 43% Superior Frontal Gyrus 41% Frontal Pole |  |
| Default mode | DMN32 | 107 | -7 | 51 | -1 | 50% Paracingulate Gyrus 38% Cingulate Gyrus, anterior division |  |
| Default mode | DMN33 | 108 | 9 | 54 | 3 | 75% Paracingulate Gyrus 13% Cingulate Gyrus, anterior division |  |
| Default mode | DMN34 | 109 | -3 | 44 | -9 | 65% Cingulate Gyrus, anterior division 32% Paracingulate Gyrus |  |
| Default mode | DMN35 | 110 | 8 | 42 | -5 | 38% Cingulate Gyrus, anterior division 11% Paracingulate Gyrus |  |
| Default mode | DMN36 | 111 | # | 45 | 8 | 30% Cingulate Gyrus, anterior division 16% Paracingulate Gyrus |  |
| Default mode | DMN37 | 112 | -2 | 38 | 36 | 69% Paracingulate Gyrus 23% Superior Frontal Gyrus |  |
| Default mode | DMN38 | 113 | -3 | 42 | 16 | 61% Cingulate Gyrus, anterior division 35% Paracingulate Gyrus |  |
| Default mode | DMN39 | 114 | # | 64 | 19 | 78% Frontal Pole |  |
| Default mode | DMN4 | 77 | # | # | 1 | 24% Cingulate Gyrus, posterior division 10% Parahippocampal Gyrus, posterior division | 35% Left Hippocampus |
| Default mode | DMN40 | 115 | -8 | 48 | 23 | 53% Paracingulate Gyrus 16% Superior Frontal Gyrus |  |
| Default mode | DMN41 | 116 | 65 | # | # | 55% Middle Temporal Gyrus, posterior division 9% Middle Temporal Gyrus, anterior division |  |
| Default mode | DMN42 | 117 | # | # | # | 28% Middle Temporal Gyrus, posterior division 17% Superior Temporal Gyrus, posterior division 12% Middle Temporal Gyrus, anterior division |  |
| Default mode | DMN43 | 118 | # | # | -4 | 50% Middle Temporal Gyrus, posterior division 14% Superior Temporal Gyrus, posterior division |  |
| Default mode | DMN44 | 119 | 65 | # | -9 | 36% Middle Temporal Gyrus, posterior division 7% Middle Temporal Gyrus, temporooccipital part |  |
| Default mode | DMN45 | 120 | # | # | -5 | 54% Middle Temporal Gyrus, posterior division 29% Middle Temporal Gyrus, temporooccipital part |  |
| Default mode | DMN46 | 121 | 13 | 30 | 59 | 63% Superior Frontal Gyrus |  |
| Default mode | DMN47 | 122 | 12 | 36 | 20 | 54% Cingulate Gyrus, anterior division 17% Paracingulate Gyrus |  |
| Default mode | DMN48 | 123 | 52 | -2 | # | 16% Superior Temporal Gyrus, anterior division 12% Superior Temporal Gyrus, posterior division |  |
| Default mode | DMN49 | 124 | # | # | -8 | 26% Lingual Gyrus 18% Parahippocampal Gyrus, posterior division | 7% Left Hippocampus |
| Default mode | DMN5 | 78 | # | 63 | -9 | 22% Frontal Pole |  |
| Default mode | DMN50 | 125 | 27 | # | # | 47% Lingual Gyrus 21% Parahippocampal Gyrus, posterior division 14% Temporal Occipital Fusiform Cortex 8% Temporal Fusiform Cortex, posterior division |  |
| Default mode | DMN51 | 126 | # | # | # | 58% Temporal Fusiform Cortex, posterior division 16% Parahippocampal Gyrus, posterior division |  |
| Default mode | DMN52 | 127 | 28 | # | # |  |  |
| Default mode | DMN53 | 128 | 52 | 7 | # | 29% Middle Temporal Gyrus, anterior division 21% Superior Temporal Gyrus, anterior division 11% Temporal Pole |  |
| Default mode | DMN54 | 129 | # | 3 | # | 33% Middle Temporal Gyrus, anterior division 11% Superior Temporal Gyrus, anterior division |  |
| Default mode | DMN55 | 130 | 47 | # | 29 | 49% Angular Gyrus |  |
| Default mode | DMN56 | 131 | # | # | 1 | 18% Middle Temporal Gyrus, posterior division 6% Middle Temporal Gyrus, temporooccipital part |  |
| Default mode | DMN57 | 137 | # | 31 | # | 59% Frontal Orbital Cortex 5% Inferior Frontal Gyrus, pars triangularis |  |
| Default mode | DMN58 | 139 | 49 | 35 | # | 32% Frontal Orbital Cortex 20% Inferior Frontal Gyrus, pars triangularis |  |
| Default mode | DMN6 | 79 | # | # | 21 | 29% Angular Gyrus 28% Lateral Occipital Cortex, superior division 7% Lateral Occipital Cortex, inferior division 5% Middle Temporal Gyrus, temporooccipital part |  |
| Default mode | DMN7 | 80 | 43 | # | 28 | 57% Lateral Occipital Cortex, superior division |  |
| Default mode | DMN8 | 81 | # | 12 | # | 19% Temporal Pole |  |
| Default mode | DMN9 | 82 | 46 | 16 | # | 50% Temporal Pole |  |
| Dorsal attention | DAN1 | 251 | 10 | # | 61 | 51% Precuneous Cortex 15% Lateral Occipital Cortex, superior division |  |
| Dorsal attention | DAN10 | 263 | # | # | 64 | 30% Lateral Occipital Cortex, superior division 19% Superior Parietal Lobule |  |
| Dorsal attention | DAN11 | 264 | 29 | -5 | 54 | 31% Precentral Gyrus 19% Superior Frontal Gyrus 5% Middle Frontal Gyrus |  |
| Dorsal attention | DAN2 | 252 | # | # | 5 | 31% Lateral Occipital Cortex, inferior division 25% Middle Temporal Gyrus, temporooccipital part |  |
| Dorsal attention | DAN3 | 256 | 22 | # | 48 | 31% Lateral Occipital Cortex, superior division 5% Precuneous Cortex |  |
| Dorsal attention | DAN4 | 257 | 46 | # | 4 | 27% Middle Temporal Gyrus, temporooccipital part 23% Lateral Occipital Cortex, inferior division |  |
| Dorsal attention | DAN5 | 258 | 25 | # | 60 | 35% Lateral Occipital Cortex, superior division |  |
| Dorsal attention | DAN6 | 259 | # | # | 47 | 38% Superior Parietal Lobule 22% Supramarginal Gyrus, posterior division 5% Angular Gyrus |  |
| Dorsal attention | DAN7 | 260 | # | # | 37 | 65% Lateral Occipital Cortex, superior division |  |
| Dorsal attention | DAN8 | 261 | # | -1 | 54 | 34% Precentral Gyrus 32% Middle Frontal Gyrus |  |
| Dorsal attention | DAN9 | 262 | # | # | -9 | 28% Temporal Occipital Fusiform Cortex 13% Inferior Temporal Gyrus, temporooccipital part 9% Occipital Fusiform Gyrus |  |
| Fronto-parietal Task Control | FPN1 | 174 | # | 2 | 46 | 42% Precentral Gyrus 14% Middle Frontal Gyrus |  |
| Fronto-parietal Task Control | FPN10 | 187 | # | 6 | 33 | 37% Precentral Gyrus 19% Middle Frontal Gyrus |  |
| Fronto-parietal Task Control | FPN11 | 188 | # | 38 | 21 | 48% Middle Frontal Gyrus 8% Frontal Pole |  |
| Fronto-parietal Task Control | FPN12 | 189 | 38 | 43 | 15 | 20% Frontal Pole 5% Middle Frontal Gyrus |  |
| Fronto-parietal Task Control | FPN13 | 190 | 49 | # | 45 | 36% Supramarginal Gyrus, posterior division 20% Angular Gyrus |  |
| Fronto-parietal Task Control | FPN14 | 191 | # | # | 48 | 29% Lateral Occipital Cortex, superior division 21% Superior Parietal Lobule 9% Angular Gyrus |  |
| Fronto-parietal Task Control | FPN15 | 192 | 44 | # | 47 | 37% Angular Gyrus 14% Lateral Occipital Cortex, superior division 11% Superior Parietal Lobule |  |
| Fronto-parietal Task Control | FPN16 | 193 | 32 | 14 | 56 | 43% Middle Frontal Gyrus 22% Superior Frontal Gyrus |  |
| Fronto-parietal Task Control | FPN17 | 194 | 37 | # | 40 | 50% Lateral Occipital Cortex, superior division |  |
| Fronto-parietal Task Control | FPN18 | 195 | # | # | 45 | 28% Angular Gyrus 11% Supramarginal Gyrus, posterior division 8% Lateral Occipital Cortex, superior division 5% Superior Parietal Lobule |  |
| Fronto-parietal Task Control | FPN19 | 196 | 40 | 18 | 40 | 39% Middle Frontal Gyrus |  |
| Fronto-parietal Task Control | FPN2 | 175 | 48 | 25 | 27 | 42% Middle Frontal Gyrus 21% Inferior Frontal Gyrus, pars opercularis 5% Inferior Frontal Gyrus, pars triangularis |  |
| Fronto-parietal Task Control | FPN20 | 197 | # | 55 | 4 | 56% Frontal Pole |  |
| Fronto-parietal Task Control | FPN21 | 198 | # | 45 | -2 | 41% Frontal Pole |  |
| Fronto-parietal Task Control | FPN22 | 199 | 33 | # | 44 | 28% Superior Parietal Lobule 11% Angular Gyrus 10% Lateral Occipital Cortex, superior division |  |
| Fronto-parietal Task Control | FPN23 | 200 | 43 | 49 | -2 | 65% Frontal Pole |  |
| Fronto-parietal Task Control | FPN24 | 201 | # | 25 | 30 | 44% Middle Frontal Gyrus 6% Inferior Frontal Gyrus, pars triangularis 5% Inferior Frontal Gyrus, pars opercularis |  |
| Fronto-parietal Task Control | FPN25 | 202 | -3 | 26 | 44 | 71% Paracingulate Gyrus 18% Superior Frontal Gyrus |  |
| Fronto-parietal Task Control | FPN3 | 176 | # | 11 | 23 | 30% Precentral Gyrus 29% Inferior Frontal Gyrus, pars opercularis |  |
| Fronto-parietal Task Control | FPN4 | 177 | # | # | 43 | 34% Supramarginal Gyrus, posterior division 21% Angular Gyrus |  |
| Fronto-parietal Task Control | FPN5 | 178 | # | 11 | 64 | 41% Superior Frontal Gyrus 5% Middle Frontal Gyrus |  |
| Fronto-parietal Task Control | FPN6 | 179 | 58 | # | # | 62% Inferior Temporal Gyrus, temporooccipital part 20% Middle Temporal Gyrus, temporooccipital part |  |
| Fronto-parietal Task Control | FPN7 | 180 | 24 | 45 | # | 21% Frontal Pole 7% Frontal Orbital Cortex |  |
| Fronto-parietal Task Control | FPN8 | 181 | 34 | 54 | # | 10% Frontal Pole |  |
| Fronto-parietal Task Control | FPN9 | 186 | 47 | 10 | 33 | 39% Precentral Gyrus 23% Middle Frontal Gyrus |  |
| Salience | SN1 | 203 | 11 | # | 50 | 56% Precuneous Cortex 22% Cingulate Gyrus, posterior division 7% Postcentral Gyrus |  |
| Salience | SN10 | 212 | # | 26 | 25 | 30% Cingulate Gyrus, anterior division 7% Paracingulate Gyrus |  |
| Salience | SN11 | 213 | -1 | 15 | 44 | 59% Paracingulate Gyrus 21% Cingulate Gyrus, anterior division 11% Juxtapositional Lobule Cortex (formerly Supplementary Motor Cortex) |  |
| Salience | SN12 | 214 | # | 52 | 21 | 78% Frontal Pole |  |
| Salience | SN13 | 215 | 0 | 30 | 27 | 68% Cingulate Gyrus, anterior division 24% Paracingulate Gyrus |  |
| Salience | SN14 | 216 | 5 | 23 | 37 | 58% Paracingulate Gyrus 37% Cingulate Gyrus, anterior division |  |
| Salience | SN15 | 217 | 10 | 22 | 27 | 36% Cingulate Gyrus, anterior division 5% Paracingulate Gyrus |  |
| Salience | SN16 | 218 | 31 | 56 | 14 | 75% Frontal Pole |  |
| Salience | SN17 | 219 | 26 | 50 | 27 | 62% Frontal Pole 5% Superior Frontal Gyrus |  |
| Salience | SN18 | 220 | # | 51 | 17 | 79% Frontal Pole |  |
| Salience | SN2 | 204 | 55 | # | 37 | 39% Angular Gyrus 25% Supramarginal Gyrus, posterior division |  |
| Salience | SN3 | 205 | 42 | 0 | 47 | 34% Precentral Gyrus 17% Middle Frontal Gyrus |  |
| Salience | SN4 | 206 | 31 | 33 | 26 | 17% Middle Frontal Gyrus |  |
| Salience | SN5 | 207 | 48 | 22 | 10 | 19% Inferior Frontal Gyrus, pars opercularis |  |
| Salience | SN6 | 208 | # | 20 | 0 | 74% Insular Cortex 5% Frontal Operculum Cortex |  |
| Salience | SN7 | 209 | 36 | 22 | 3 | 57% Insular Cortex 21% Frontal Operculum Cortex |  |
| Salience | SN8 | 210 | 37 | 32 | -2 | 29% Frontal Operculum Cortex 25% Frontal Orbital Cortex 11% Insular Cortex 10% Inferior Frontal Gyrus, pars triangularis |  |
| Salience | SN9 | 211 | 34 | 16 | -8 | 10% Insular Cortex |  |
| Subcortical | SC1 | 222 | 6 | # | 0 |  | 55% Right Thalamus |
| Subcortical | SC10 | 231 | 29 | 1 | 4 |  | 100% Right Putamen |
| Subcortical | SC11 | 232 | # | # | 0 |  | 98% Left Putamen |
| Subcortical | SC12 | 233 | 15 | 5 | 7 |  | 87% 7% Right Thalamus 6% Right Caudate |
| Subcortical | SC13 | 234 | 9 | -4 | 6 |  | 100% Right Thalamus |
| Subcortical | SC2 | 223 | -2 | # | 12 |  | 68% Left Thalamus |
| Subcortical | SC3 | 224 | # | # | 7 |  | 68% Left Thalamus |
| Subcortical | SC4 | 225 | 12 | # | 8 |  | 68% Left Thalamus |
| Subcortical | SC5 | 226 | -5 | # | -4 |  | 70% Brain-Stem |
| Subcortical | SC6 | 227 | # | 7 | -5 |  | 82% Left Putamen 18% Left Pallidum |
| Subcortical | SC7 | 228 | # | 4 | 8 |  |  |
| Subcortical | SC8 | 229 | 31 | # | 2 |  | 90% Right Putamen 8% |
| Subcortical | SC9 | 230 | 23 | 10 | 1 |  | 94% Right Putamen 6% |
| Ventral attention | VAN1 | 138 | # | 11 | 67 | 53% Superior Frontal Gyrus |  |
| Ventral attention | VAN2 | 235 | 54 | # | 22 | 25% Angular Gyrus 20% Supramarginal Gyrus, posterior division |  |
| Ventral attention | VAN3 | 236 | # | # | 10 | 47% Middle Temporal Gyrus, temporooccipital part 15% Supramarginal Gyrus, posterior division 14% Angular Gyrus |  |
| Ventral attention | VAN4 | 237 | # | # | 14 | 15% Supramarginal Gyrus, posterior division 9% Superior Temporal Gyrus, posterior division 5% Planum Temporale |  |
| Ventral attention | VAN5 | 238 | 52 | # | 8 | 16% Supramarginal Gyrus, posterior division 10% Superior Temporal Gyrus, posterior division |  |
| Ventral attention | VAN6 | 239 | 51 | # | -4 | 36% Middle Temporal Gyrus, posterior division 23% Superior Temporal Gyrus, posterior division 5% Supramarginal Gyrus, posterior division |  |
| Ventral attention | VAN7 | 240 | 56 | # | 11 | 35% Middle Temporal Gyrus, temporooccipital part 22% Angular Gyrus 9% Supramarginal Gyrus, posterior division |  |
| Ventral attention | VAN8 | 241 | 53 | 33 | 1 | 34% Inferior Frontal Gyrus, pars triangularis 5% Inferior Frontal Gyrus, pars opercularis |  |
| Ventral attention | VAN9 | 242 | # | 25 | -1 | 32% Inferior Frontal Gyrus, pars triangularis 27% Frontal Operculum Cortex 14% Inferior Frontal Gyrus, pars opercularis |  |

*Supplemental Table 7: D-MRI ROI abbreviations:*

| Full tract name | Abbreviation |
| --- | --- |
| Full skeleton average FA | Average FA |
| Anterior corona radiata | ACR (L+R) |
| Anterior limb of internal capsule | ALIC (L+R) |
| Body of corpus callosum | BCC |
| Cingulum (cingulate gyrus) | CGC (L+R) |
| Cingulum (hippocampal portion) | CGH (L+R) |
| Corona radiata | CR (L+R) |
| External capsule | EC (L+R) |
| Fornix | FX |
| Fornix (crus) / Stria terminalis | FXST (L+R) |
| Genu of corpus callosum | GCC |
| Internal capsule | IC (L+R) |
| Posterior corona radiata | PCR (L+R) |
| Posterior limb of internal capsule | PLIC (L+R) |
| Posterior thalamic radiation | PTR (L+R) |
| Retrolenticular part of internal capsule | RLIC (L+R) |
| Splenium of corpus callosum | SCC |
| Superior corona radiata | SCR (L+R) |
| Superior fronto-occipital fasciculus | SFO (L+R) |
| Superior longitudinal fasciculus | SLF (L+R) |
| Sagittal stratum | SS (L+R) |
| Corticospinal tract | CST (L+R) |
| Uncinate fasciculus | UNC (L+R) |
| Tapetum | TAP (L+R) |

*Supplemental Table 8: ML performance for each modality*

| **Modality** | | **T1** | | | **RS** | | | **DTI** | | |
| --- | --- | --- | --- | --- | --- | --- | --- | --- | --- | --- |
| **Control type** | | Control | HC | TEHC | Control | HC | TEHC | Control | HC | TEHC |
| **N Features** | | 97 | 97 | 97 | 10878 | 10878 | 10878 | 40 | 40 | 40 |
| **RF** | **CV Accuracy** | 0.62 | 0.65 | 0.58 | 0.59 | 0.77 | 0.64 | 0.55 | 0.65 | 0.66 |
|  | **CV AUC** | 0.59 | 0.72 | 0.6 | 0.55 | 0.7 | 0.64 | 0.55 | 0.74 | 0.59 |
|  | **test Sensitivity** | 0.56 | 0.74 | 0.51 | 0.49 | 0.68 | 0.62 | 0.53 | 0.75 | 0.6 |
|  | **test Specificity** | 0.64 | 0.6 | 0.7 | 0.59 | 0.64 | 0.58 | 0.59 | 0.68 | 0.51 |
|  | **test Accuracy** | 0.59 | 0.75 | 0.6 | 0.55 | 0.66 | 0.57 | 0.56 | 0.68 | 0.55 |
|  | **test AUC** | **0.61** | **0.73** | **0.64** | **0.58** | **0.72** | **0.63** | **0.59** | **0.74** | **0.59** |
| **SVM** | **CV Accuracy** | 0.56 | 0.68 | 0.57 | 0.59 | 0.75 | 0.64 | 0.53 | 0.76 | 0.61 |
|  | **CV AUC** | 0.6 | 0.73 | 0.6 | 0.6 | 0.77 | 0.68 | 0.51 | 0.79 | 0.61 |
|  | **test Sensitivity** | 0.56 | 0.8 | 0.61 | 0.49 | 0.69 | 0.6 | 0.63 | 0.68 | 0.6 |
|  | **test Specificity** | 0.56 | 0.58 | 0.6 | 0.66 | 0.68 | 0.67 | 0.57 | 0.75 | 0.55 |
|  | **test Accuracy** | 0.56 | 0.74 | 0.6 | 0.58 | 0.7 | 0.64 | 0.47 | 0.68 | 0.57 |
|  | **test AUC** | **0.60** | **0.72** | **0.64** | **0.59** | **0.75** | **0.69** | **0.56** | **0.78** | **0.59** |

*Supplemental Table 9: ML performance for each modality by site*

|  | Site | N_PTSD | N_Control | RF | | | | SVM | | | |
| --- | --- | --- | --- | --- | --- | --- | --- | --- | --- | --- | --- |
|  |  |  |  | **Accuracy** | **AUC** | **Sensitivity** | **Specificity** | **Accuracy** | **AUC** | **Sensitivity** | **Specificity** |
| s-MRI | Uwash | 54 | 192 | 0.51 | 0.55 | 0.6 | 0.58 | 0.52 | 0.55 | **0.58** | **0.54** |
|  | ADNIDOD | 80 | 114 | 0.5 | 0.56 | 0.59 | 0.45 | 0.52 | 0.54 | 0.44 | 0.59 |
|  | AMC | 38 | 37 | 0.47 | 0.52 | 0.44 | 0.45 | 0.5 | 0.47 | 0.56 | 0.41 |
|  | Bruce | 66 | 19 | 0.44 | 0.47 | 0.4 | 0.43 | 0.45 | 0.43 | 0.45 | 0.43 |
|  | Columbia | 53 | 35 | 0.47 | 0.47 | 0.46 | 0.54 | 0.53 | 0.53 | 0.58 | 0.49 |
|  | DUKE | 112 | 264 | 0.53 | 0.52 | 0.54 | 0.5 | 0.51 | 0.53 | 0.49 | 0.52 |
|  | DUKE_DeBellis | 30 | 88 | 0.59 | 0.61 | 0.56 | 0.63 | 0.57 | 0.55 | 0.52 | 0.55 |
|  | Grupe | 19 | 39 | 0.58 | 0.6 | 0.53 | 0.6 | 0.55 | 0.6 | 0.43 | 0.49 |
|  | GTP | 59 | 115 | 0.54 | 0.5 | 0.52 | 0.51 | 0.48 | 0.49 | 0.44 | 0.52 |
|  | intrust | 108 | 258 | 0.54 | 0.57 | 0.53 | 0.54 | 0.54 | 0.59 | 0.62 | 0.55 |
|  | Leiden | 22 | 30 | 0.46 | 0.42 | 0.38 | 0.44 | 0.46 | 0.44 | 0.42 | 0.56 |
|  | McLean | 39 | 13 | 0.51 | 0.6 | 0.53 | 0.58 | 0.53 | 0.53 | 0.55 | 0.48 |
|  | McLeanRosso | 22 | 91 | 0.56 | 0.55 | 0.59 | 0.56 | 0.53 | 0.62 | 0.63 | 0.5 |
|  | Munster | 21 | 26 | 0.64 | 0.72 | 0.51 | 0.69 | 0.6 | 0.63 | 0.51 | 0.71 |
|  | SouthDakota | 78 | 45 | 0.43 | 0.38 | 0.39 | 0.5 | 0.47 | 0.46 | 0.44 | 0.52 |
|  | UCAS | 34 | 36 | 0.47 | 0.51 | 0.47 | 0.55 | 0.5 | 0.51 | 0.62 | 0.41 |
|  | UCI | 13 | 15 | 0.77 | 0.78 | 0.62 | 0.67 | 0.77 | 0.83 | 0.82 | 0.67 |
|  | UIL_Ch | 24 | 20 | 0.54 | 0.67 | 0.58 | 0.5 | 0.7 | 0.83 | 0.92 | 0.4 |
|  | Umich | 26 | 43 | 0.54 | 0.5 | 0.54 | 0.55 | 0.45 | 0.47 | 0.23 | 0.56 |
|  | UNSW | 49 | 118 | 0.51 | 0.49 | 0.39 | 0.54 | 0.49 | 0.43 | 0.44 | 0.48 |
|  | Utoledo | 15 | 64 | 0.59 | 0.65 | 0.58 | 0.63 | 0.47 | 0.61 | 0.47 | 0.56 |
|  | Utrecht | 56 | 54 | 0.44 | 0.4 | 0.43 | 0.47 | 0.47 | 0.46 | 0.37 | 0.54 |
|  | VA_AA | 41 | 22 | 0.51 | 0.49 | 0.44 | 0.63 | 0.5 | 0.53 | 0.46 | 0.52 |
|  | WACO | 41 | 25 | 0.4 | 0.38 | 0.42 | 0.47 | 0.42 | 0.4 | 0.42 | 0.38 |
|  | WestHaven | 37 | 38 | 0.62 | 0.66 | 0.62 | 0.62 | 0.57 | 0.58 | 0.51 | 0.59 |
|  | WiscLarson | 20 | 47 | 0.63 | 0.72 | 0.7 | 0.65 | 0.56 | 0.68 | 0.67 | 0.55 |
|  | Yale | 23 | 49 | 0.56 | 0.61 | 0.56 | 0.55 | 0.56 | 0.58 | 0.43 | 0.63 |
|  | VAMinn | 94 | 149 | 0.55 | 0.5 | 0.49 | 0.52 | 0.53 | 0.53 | 0.48 | 0.54 |
| Rs-fMRI | AMC | 34 | 36 | 0.51 | 0.55 | 0.52 | 0.51 | 0.54 | 0.6 | 0.71 | 0.42 |
|  | Beijing | 41 | 45 | 0.55 | 0.59 | 0.52 | 0.59 | 0.56 | 0.62 | 0.64 | 0.43 |
|  | Duke | 31 | 104 | 0.5 | 0.48 | 0.39 | 0.58 | 0.57 | 0.53 | 0.37 | 0.61 |
|  | Emory | 35 | 70 | 0.48 | 0.55 | 0.51 | 0.56 | 0.51 | 0.54 | 0.51 | 0.57 |
|  | Leiden | 21 | 29 | 0.51 | 0.53 | 0.47 | 0.56 | 0.5 | 0.51 | 0.49 | 0.59 |
|  | Masaryk | 109 | 155 | 0.5 | 0.49 | 0.47 | 0.52 | 0.51 | 0.51 | 0.46 | 0.49 |
|  | McLean | 42 | 23 | 0.51 | 0.44 | 0.53 | 0.52 | 0.68 | 0.61 | 0.78 | 0.36 |
|  | VAMinn | 90 | 157 | 0.59 | 0.6 | 0.56 | 0.61 | 0.58 | 0.63 | 0.65 | 0.56 |
|  | Nanjing | 45 | 84 | 0.46 | 0.51 | 0.45 | 0.5 | 0.51 | 0.47 | 0.49 | 0.49 |
|  | Stanford | 99 | 81 | 0.55 | 0.54 | 0.52 | 0.57 | 0.62 | 0.6 | 0.65 | 0.55 |
|  | Uwash | 33 | 116 | 0.58 | 0.57 | 0.56 | 0.66 | 0.65 | 0.67 | 0.51 | 0.68 |
|  | Utrecht | 53 | 51 | 0.56 | 0.51 | 0.49 | 0.57 | 0.49 | 0.53 | 0.23 | 0.74 |
|  | WesternOntario | 106 | 52 | 0.54 | 0.55 | 0.57 | 0.51 | 0.54 | 0.61 | 0.58 | 0.45 |
| D-MRI | ADNIDOD | 69 | 66 | 0.57 | 0.58 | 0.6 | 0.49 | 0.55 | 0.56 | 0.55 | 0.49 |
|  | AMC | 34 | 36 | 0.41 | 0.38 | 0.44 | 0.45 | 0.37 | 0.33 | 0.4 | 0.33 |
|  | Beijing | 32 | 35 | 0.61 | 0.62 | 0.55 | 0.53 | 0.56 | 0.65 | 0.72 | 0.45 |
|  | Columbia | 18 | 15 | 0.41 | 0.41 | 0.33 | 0.43 | 0.32 | 0.32 | 0.45 | 0.15 |
|  | Duke | 101 | 237 | 0.52 | 0.49 | 0.47 | 0.54 | 0.65 | 0.55 | 0.36 | 0.78 |
|  | EmoryGTP | 47 | 85 | 0.44 | 0.47 | 0.45 | 0.46 | 0.53 | 0.46 | 0.37 | 0.53 |
|  | Lawson | 46 | 52 | 0.52 | 0.58 | 0.56 | 0.56 | 0.54 | 0.59 | 0.64 | 0.46 |
|  | LUMC | 20 | 20 | 0.57 | 0.6 | 0.52 | 0.58 | 0.57 | 0.63 | 0.58 | 0.57 |
|  | McLean | 41 | 14 | 0.36 | 0.32 | 0.3 | 0.37 | 0.36 | 0.35 | 0.22 | 0.62 |
|  | SouthDakota | 55 | 36 | 0.46 | 0.39 | 0.39 | 0.49 | 0.43 | 0.49 | 0.28 | 0.73 |
|  | Stellenbosch | 44 | 58 | 0.46 | 0.48 | 0.48 | 0.51 | 0.51 | 0.52 | 0.4 | 0.66 |
|  | Utrecht | 46 | 48 | 0.42 | 0.4 | 0.38 | 0.5 | 0.47 | 0.41 | 0.39 | 0.51 |
|  | VAMinn | 105 | 149 | 0.57 | 0.57 | 0.55 | 0.61 | 0.54 | 0.56 | 0.49 | 0.6 |
|  | VAWaco | 36 | 17 | 0.49 | 0.46 | 0.48 | 0.45 | 0.48 | 0.35 | 0.36 | 0.57 |
|  | GrupeWisconsin | 16 | 32 | 0.57 | 0.58 | 0.6 | 0.51 | 0.55 | 0.56 | 0.62 | 0.52 |
|  | Yale | 37 | 30 | 0.55 | 0.59 | 0.54 | 0.61 | 0.64 | 0.62 | 0.51 | 0.79 |
|  | UWash | 31 | 136 | 0.61 | 0.68 | 0.61 | 0.63 | 0.65 | 0.67 | 0.55 | 0.71 |

*Supplemental Table 10: LOSOCV ML performance for each modality*

| Modality | Site | N_PTSD | N_Control | RF | | | | | | SVM | | | | | |
| --- | --- | --- | --- | --- | --- | --- | --- | --- | --- | --- | --- | --- | --- | --- | --- |
|  |  |  |  | **CV Accuracy** | **CV AUC** | **test Sensitivity** | **test Specificity** | **test Accuracy** | **test AUC** | **CV Accuracy** | **CV AUC** | **test Sensitivity** | **test Specificity** | **test Accuracy** | **test AUC** |
| s-MRI | Uwash | 54 | 192 | 0.55 | 0.45 | 0.63 | 0.47 | 0.51 | 0.57 | 0.55 | 0.45 | 0.46 | 0.69 | 0.64 | 0.59 |
|  | ADNIDOD | 80 | 114 | 0.59 | 0.51 | 0.57 | 0.53 | 0.55 | 0.48 | 0.56 | 0.45 | 0.64 | 0.51 | 0.56 | 0.56 |
|  | AMC | 38 | 37 | 0.57 | 0.47 | 0.47 | 0.54 | 0.51 | 0.49 | 0.54 | 0.44 | 0.37 | 0.7 | 0.53 | 0.41 |
|  | Bruce | 66 | 19 | 0.54 | 0.44 | 0.58 | 0.63 | 0.59 | 0.61 | 0.54 | 0.42 | 0.85 | 0.47 | 0.76 | 0.67 |
|  | Columbia | 53 | 35 | 0.56 | 0.45 | 0.77 | 0.43 | 0.64 | 0.57 | 0.54 | 0.42 | 0.58 | 0.57 | 0.58 | 0.55 |
|  | DUKE | 112 | 264 | 0.55 | 0.48 | 0.54 | 0.52 | 0.52 | 0.54 | 0.55 | 0.46 | 0.57 | 0.48 | 0.51 | 0.52 |
|  | DUKE_DeBellis | 30 | 88 | 0.56 | 0.46 | 0.6 | 0.61 | 0.61 | 0.64 | 0.55 | 0.47 | 0.57 | 0.58 | 0.58 | 0.59 |
|  | Grupe | 19 | 39 | 0.55 | 0.46 | 0.46 | 0.6 | 0.55 | 0.51 | 0.54 | 0.44 | 0.61 | 0.51 | 0.55 | 0.52 |
|  | GTP | 59 | 115 | 0.57 | 0.46 | 0.68 | 0.59 | 0.62 | 0.63 | 0.57 | 0.46 | 0.58 | 0.33 | 0.41 | 0.43 |
|  | intrust | 108 | 258 | 0.56 | 0.47 | 0.41 | 0.77 | 0.62 | 0.62 | 0.56 | 0.44 | 0.55 | 0.67 | 0.62 | 0.56 |
|  | Leiden | 22 | 30 | 0.57 | 0.46 | 0.64 | 0.69 | 0.65 | 0.67 | 0.55 | 0.44 | 0.79 | 0.54 | 0.73 | 0.64 |
|  | McLean | 39 | 13 | 0.54 | 0.45 | 0.59 | 0.66 | 0.65 | 0.67 | 0.53 | 0.42 | 0.36 | 0.84 | 0.74 | 0.56 |
|  | McLeanRosso | 22 | 91 | 0.56 | 0.47 | 0.76 | 0.42 | 0.57 | 0.57 | 0.55 | 0.45 | 0.48 | 0.5 | 0.49 | 0.41 |
|  | Munster | 21 | 26 | 0.55 | 0.48 | 0.46 | 0.76 | 0.57 | 0.55 | 0.57 | 0.45 | 0.4 | 0.76 | 0.53 | 0.53 |
|  | SouthDakota | 78 | 45 | 0.56 | 0.46 | 0.65 | 0.64 | 0.64 | 0.61 | 0.55 | 0.45 | 0.53 | 0.53 | 0.53 | 0.52 |
|  | UCAS | 34 | 36 | 0.57 | 0.48 | 0.46 | 0.73 | 0.61 | 0.56 | 0.57 | 0.45 | 0.77 | 0.6 | 0.68 | 0.74 |
|  | UCI | 13 | 15 | 0.57 | 0.48 | 0.71 | 0.45 | 0.59 | 0.52 | 0.57 | 0.46 | 0.38 | 0.55 | 0.45 | 0.48 |
|  | UIL_Ch | 24 | 20 | 0.55 | 0.44 | 0.43 | 0.67 | 0.6 | 0.52 | 0.55 | 0.43 | 0.43 | 0.58 | 0.54 | 0.48 |
|  | Umich | 26 | 43 | 0.56 | 0.46 | 0.81 | 0.35 | 0.52 | 0.54 | 0.55 | 0.45 | 0.54 | 0.35 | 0.42 | 0.42 |
|  | UNSW | 49 | 118 | 0.55 | 0.44 | 0.73 | 0.56 | 0.59 | 0.7 | 0.55 | 0.43 | 0.47 | 0.8 | 0.73 | 0.65 |
|  | Utoledo | 15 | 64 | 0.57 | 0.48 | 0.57 | 0.61 | 0.59 | 0.58 | 0.58 | 0.46 | 0.5 | 0.67 | 0.58 | 0.58 |
|  | Utrecht | 56 | 54 | 0.55 | 0.45 | 0.47 | 0.54 | 0.51 | 0.49 | 0.54 | 0.41 | 0.57 | 0.5 | 0.53 | 0.54 |
|  | VA_AA | 41 | 22 | 0.57 | 0.49 | 0.63 | 0.45 | 0.57 | 0.5 | 0.59 | 0.47 | 0.41 | 0.68 | 0.51 | 0.53 |
|  | WACO | 41 | 25 | 0.58 | 0.48 | 0.37 | 0.56 | 0.44 | 0.46 | 0.58 | 0.47 | 0.46 | 0.68 | 0.55 | 0.57 |
|  | WestHaven | 37 | 38 | 0.57 | 0.49 | 0.49 | 0.68 | 0.59 | 0.58 | 0.57 | 0.46 | 0.68 | 0.53 | 0.6 | 0.61 |
|  | WiscLarson | 20 | 47 | 0.55 | 0.46 | 0.4 | 0.51 | 0.48 | 0.41 | 0.55 | 0.44 | 0.55 | 0.72 | 0.67 | 0.67 |
|  | Yale | 23 | 49 | 0.56 | 0.46 | 0.61 | 0.71 | 0.68 | 0.65 | 0.56 | 0.45 | 0.48 | 0.71 | 0.64 | 0.57 |
|  | VAMinn | 94 | 149 | 0.56 | 0.47 | 0.61 | 0.52 | 0.54 | 0.58 | 0.55 | 0.46 | 0.56 | 0.49 | 0.51 | 0.51 |
| RS-fMRI | AMC | 34 | 36 | 0.54 | 0.45 | 0.71 | 0.39 | 0.54 | 0.55 | 0.52 | 0.4 | 0.74 | 0.36 | 0.54 | 0.52 |
|  | Beijing | 41 | 45 | 0.53 | 0.44 | 0.51 | 0.58 | 0.55 | 0.52 | 0.51 | 0.39 | 0.54 | 0.56 | 0.55 | 0.52 |
|  | Duke | 31 | 104 | 0.55 | 0.54 | 0.61 | 0.36 | 0.41 | 0.44 | 0.54 | 0.51 | 0.48 | 0.37 | 0.39 | 0.37 |
|  | Emory | 35 | 70 | 0.53 | 0.48 | 0.34 | 0.67 | 0.56 | 0.5 | 0.52 | 0.45 | 0.69 | 0.39 | 0.49 | 0.52 |
|  | Leiden | 21 | 29 | 0.52 | 0.44 | 0.43 | 0.55 | 0.5 | 0.43 | 0.52 | 0.42 | 0.57 | 0.48 | 0.52 | 0.42 |
|  | Masaryk | 109 | 155 | 0.58 | 0.54 | 0.49 | 0.52 | 0.5 | 0.46 | 0.57 | 0.53 | 0.53 | 0.43 | 0.47 | 0.46 |
|  | McLean | 42 | 23 | 0.55 | 0.44 | 0.6 | 0.59 | 0.59 | 0.55 | 0.53 | 0.4 | 0.55 | 0.54 | 0.55 | 0.49 |
|  | VAMinn | 90 | 157 | 0.52 | 0.47 | 0.77 | 0.42 | 0.55 | 0.59 | 0.51 | 0.43 | 0.56 | 0.56 | 0.56 | 0.58 |
|  | Nanjing | 45 | 84 | 0.53 | 0.48 | 0.47 | 0.67 | 0.6 | 0.58 | 0.51 | 0.43 | 0.44 | 0.65 | 0.58 | 0.51 |
|  | Stanford | 99 | 81 | 0.56 | 0.54 | 0.38 | 0.79 | 0.57 | 0.56 | 0.55 | 0.48 | 0.37 | 0.78 | 0.56 | 0.54 |
|  | Uwash | 33 | 116 | 0.47 | 0.39 | 0.45 | 0.59 | 0.56 | 0.52 | 0.42 | 0.31 | 0.48 | 0.47 | 0.48 | 0.5 |
|  | Utrecht | 53 | 51 | 0.54 | 0.49 | 0.62 | 0.57 | 0.6 | 0.6 | 0.53 | 0.48 | 0.7 | 0.41 | 0.56 | 0.52 |
|  | WesternOntario | 106 | 52 | 0.57 | 0.49 | 0.79 | 0.38 | 0.66 | 0.58 | 0.59 | 0.46 | 0.57 | 0.54 | 0.56 | 0.54 |
| D-MRI | ADNIDOD | 69 | 66 | 0.42 | 0.34 | 0.48 | 0.7 | 0.59 | 0.54 | 0.48 | 0.33 | 0.29 | 0.74 | 0.51 | 0.47 |
|  | AMC | 34 | 36 | 0.42 | 0.33 | 0.35 | 0.58 | 0.47 | 0.45 | 0.48 | 0.34 | 0.65 | 0.5 | 0.57 | 0.55 |
|  | Beijing | 32 | 35 | 0.44 | 0.35 | 0.47 | 0.74 | 0.61 | 0.62 | 0.49 | 0.37 | 0.69 | 0.51 | 0.6 | 0.61 |
|  | Columbia | 18 | 15 | 0.45 | 0.37 | 0.56 | 0.53 | 0.55 | 0.54 | 0.5 | 0.38 | 0.67 | 0.53 | 0.61 | 0.59 |
|  | Duke | 101 | 237 | 0.38 | 0.32 | 0.51 | 0.59 | 0.57 | 0.55 | 0.42 | 0.32 | 0.5 | 0.61 | 0.57 | 0.57 |
|  | EmoryGTP | 47 | 85 | 0.41 | 0.37 | 0.43 | 0.59 | 0.53 | 0.51 | 0.47 | 0.37 | 0.47 | 0.48 | 0.48 | 0.47 |
|  | Lawson | 46 | 52 | 0.45 | 0.38 | 0.31 | 0.66 | 0.54 | 0.43 | 0.51 | 0.4 | 0.56 | 0.5 | 0.52 | 0.53 |
|  | LUMC | 20 | 20 | 0.46 | 0.4 | 0.55 | 0.75 | 0.65 | 0.6 | 0.51 | 0.4 | 0.35 | 0.5 | 0.42 | 0.38 |
|  | McLean | 41 | 14 | 0.44 | 0.37 | 0.52 | 0.56 | 0.54 | 0.54 | 0.49 | 0.38 | 0.54 | 0.52 | 0.53 | 0.52 |
|  | SouthDakota | 55 | 36 | 0.48 | 0.41 | 0.56 | 0.57 | 0.56 | 0.58 | 0.54 | 0.42 | 0.44 | 0.64 | 0.49 | 0.53 |
|  | Stellenbosch | 44 | 58 | 0.47 | 0.41 | 0.47 | 0.5 | 0.48 | 0.47 | 0.54 | 0.42 | 0.45 | 0.53 | 0.48 | 0.46 |
|  | Utrecht | 46 | 48 | 0.43 | 0.37 | 0.3 | 0.76 | 0.56 | 0.43 | 0.49 | 0.38 | 0.23 | 0.78 | 0.54 | 0.36 |
|  | VAMinn | 105 | 149 | 0.43 | 0.37 | 0.74 | 0.53 | 0.57 | 0.61 | 0.45 | 0.39 | 0.39 | 0.49 | 0.47 | 0.37 |
|  | VAWaco | 36 | 17 | 0.46 | 0.39 | 0.33 | 0.58 | 0.46 | 0.42 | 0.5 | 0.38 | 0.52 | 0.67 | 0.6 | 0.53 |
|  | GrupeWisconsin | 16 | 32 | 0.44 | 0.38 | 0.45 | 0.51 | 0.48 | 0.44 | 0.47 | 0.37 | 0.55 | 0.49 | 0.52 | 0.52 |
|  | Yale | 37 | 30 | 0.44 | 0.38 | 0.61 | 0.53 | 0.58 | 0.52 | 0.53 | 0.39 | 0.42 | 0.76 | 0.53 | 0.52 |
|  | UWash | 31 | 136 | 0.46 | 0.39 | 0.54 | 0.3 | 0.43 | 0.34 | 0.53 | 0.42 | 0.38 | 0.63 | 0.49 | 0.47 |
